# Influenza evolution/adaptation samples a highly non-random error landscape for hemagglutinin-encoding RNA

**DOI:** 10.64898/2026.06.15.732443

**Authors:** Olga Barranco-Gómez, Margarita Barriga, Alberto Fernandez-Fernandez, Laura García-Corzo, Antón Vizcaíno, Paula Ramilo, Antonio Osuna, Valeria A. Risso, Jose M. Sanchez-Ruiz

## Abstract

Natural selection acts on diversity generated by errors in the biosynthesis of the genetic material. Previous work has shown, however, that such errors are not necessarily fully random. We have used a model influenza strain and a unique-molecular-identifier-based high-throughput sequencing approach to assess the error landscape for hemagglutinin-encoding RNA. Single-site errors occur at highly variable frequencies, with differences that span several orders of magnitude, plausibly reflecting specific RNA sequence/structure patterns. Remarkably, influenza evolution/adaptation preferentially selects mutations encoded by the higher frequency errors, as shown by analyses of mutations fixed in natural strains over many decades and by analyses of antibody-escape mutations found in laboratory experiments on strains of the 2009 pandemics. Our results support that RNA error landscapes may provide information useful for predicting influenza evolution and point to high-frequency errors encoding antibody-evading mutations as potential contributors to the rapid evolution of influenza viruses.

## INTRODUCTION

Influenza vaccines show limited effectiveness (Zimmerman et al., 2016; Rolfes et al., 2019), a fact commonly attributed to the rapid evolution of influenza viruses and the relatively long vaccine production timescale (Gerdil, 2003; Łuksza and Lässig, 2014; Shi et al., 2025). Anticipating future evolution of influenza viruses is, therefore, of considerable interest. Efforts have focused on predicting amino acid sequence evolution for hemagglutinin (Shi et al., 2025, and references quoted therein), the viral protein responsible for the primary interaction with the host cell surface and a main target for neutralizing antibodies.

Natural selection acts on diversity generated by errors in the biosynthesis of the genetic material. Such errors, however, are not necessarily fully random (Acevedo et al., 2014; Gout et al., 2017). In fact, RNA sequence/structure patterns may promote specific and even “useful” errors, as illustrated by the fact that some viruses use programmed recoding (frameshifting and stop-codon misreading) to generate functional protein variants at significant levels (Brierley et al., 2010; Miller and Griedoc, 2010; Namy and Russel, 2010). Here, we provide evidence that the single-site error landscape for hemagglutinin-encoding RNAs is highly non-random and that amino acid replacements that are fixed during influenza evolution/adaptation are preferentially encoded by the higher frequency errors.

We have determined the single-site error landscapes for the two hemagglutinin-encoding RNAs from the A/WSN/1933 H1N1 influenza strain: the viral (virion encapsulated) RNA (vRNA) and the messenger RNA (mRNA). Since errors occur at a low level, we have used a unique molecular identifier (UMI) approach in high-throughput sequencing (Kinde et al., 2011) to set actual errors apart from common sequencing errors. We find that the single-site errors (single-nucleotide substitutions) occur at highly variable frequencies, with differences that span several orders of magnitude, plausibly reflecting specific RNA sequence/structure patterns. Adaptation is expected to occur at the amino acid sequence level. We show that the highly non-random RNA error landscape translates into a highly non-random landscape for amino acid replacements.

The availability of experimental error landscapes allows us to assess the extent to which influenza evolution/adaptation is determined by non-random diversity in the genetic material. Our analyses reveal a significant statistical correspondence between fixed mutations in the hemagglutinin gene and high-frequency errors. Specifically, we find preferential selection of the higher frequencies in the error landscape, i) when considering mutations fixed in natural influenza strains over many decades, and ii) when considering escape mutations found in laboratory experiments on antibody evasion with H1 strains of the 2009 pandemics.

According to our results and analyses, the likelihood that a given mutation is fixed depends, not only on its capability to disrupt antibody binding while maintaining viral fitness, but also on the frequency of its encoding error. It follows that RNA error landscapes may provide information useful for predicting influenza evolution.

Our results and analyses suggest a mechanism potentially contributing to the rapid evolution of influenza viruses. Antibody-evading mutations encoded by high-frequency errors will display enhanced fixation probability, thus promoting fast adaptation to the host immune response. The existence of this mechanism could have an impact in vaccine design.

## RESULTS AND DISCUSSION

### SafeSeqS determination of RNA error landscapes

Assessing sequence error landscapes is challenging, because the error rate of standard high-throughput sequencing methodologies is too high for applications that require the detection of low-frequency variants with high confidence. Here, we have resorted to the SafeSeqS methodology, which was originally developed for DNA sequencing (Kinde et al., 2011). SafeSeqS relies on the assignment of a unique molecular identifier (UMI) to each DNA template. Upon amplification, an UMI family of identical sequences is generated from each tagged template. Unlike most sequencing errors, a single-site variation actually present in the original template pool will appear in all (or, at least most) of the sequences in one or several of the resulting families.

Briefly (see Methods for details), we designed a set of four primers spanning the whole hemagglutinin genetic sequence in fragments of 400-500 bp each (Table S1). The primers included a 12 bp random sequence at the 5’ end to serve as UMI in the SafeSeqS procedure. Uniquely UMI-tagged molecules were amplified by PCR to generate multiple copies for each UMI (*i.e.*, to generate UMI families). The pool was sequenced in a fraction of a NovaSeq flow cell with a PE250 chemistry (Illumina) aiming for a total output of 12 gigabases. The UMI family distributions (Table S2, Figures S1-S4) were exponential-like, with most of the retained families having two members, but also with a substantial number of families having three or more members. Following the original work (Kinde et al., 2011), a family is taken to identify an actual substitution if it appears in at least 95% of the family members (see Methods for details). Frequencies for the detected single-site changes in the template pool were calculated as the ratio of the number of families bearing the change over the total number of families. Frequencies could be calculated for all the possible 5085 single-site changes for an RNA of length 1695 nucleotides (Tables S3-S6), although some nucleotide changes were not observed at certain positions.

It is very important to note that, while the SafeSeqS methodology was originally developed for DNA sequencing (Kinde et al., 2011), it is applied in this work to RNA sequencing. Therefore, a reverse transcription step is required to obtain a cDNA copy of the RNA. This reverse transcription is performed using one of the UMI-tagged primers. This is followed by a one-cycle PCR that incorporates the UMI at the other end of the molecule. Once both UMIs are attached, a 30-cycle PCR is performed in order to amplify the DNA. Errors during the reverse transcription step or during the one-cycle PCR would propagate through all family members, making them indistinguishable from the actual errors in RNA biosynthesis we seek to determine. On the other hand, errors introduced in the 30-cycle PCR will be discarded in the downstream bioinformatics analyses and are not expected to significantly affect the results. The impact of the reverse transcription and one-cycle PCR steps on the determined error landscapes is discussed further below.

### Error landscape for hemagglutinin-encoding RNAs

We have applied the SafeSeqS methodology, as described in the preceding section, to hemagglutinin-encoding messenger RNA (mRNA) and viral (*i.e.,* virion-encapsulated) RNA (vRNA) from the A/WSN/1933 H1N1influenza strain (Tables S3-S6 and Figure 1). Note that A/WSN/1933 (Stuart-Harris and Lond, 1939), a widely used model strain, is a laboratory adaptation of the original A/Wilson-Smith/1933 strain (Smith et al., 1933), which is closely related to the 1918 flu (Thyagarajan and Bloom, 2014). There are only 7 nucleotide differences between the hemagglutinin-encoding RNA sequences from the A/WSN/1933 and the A/Wilson-Smith/1933 strains.

**Figure 1.**
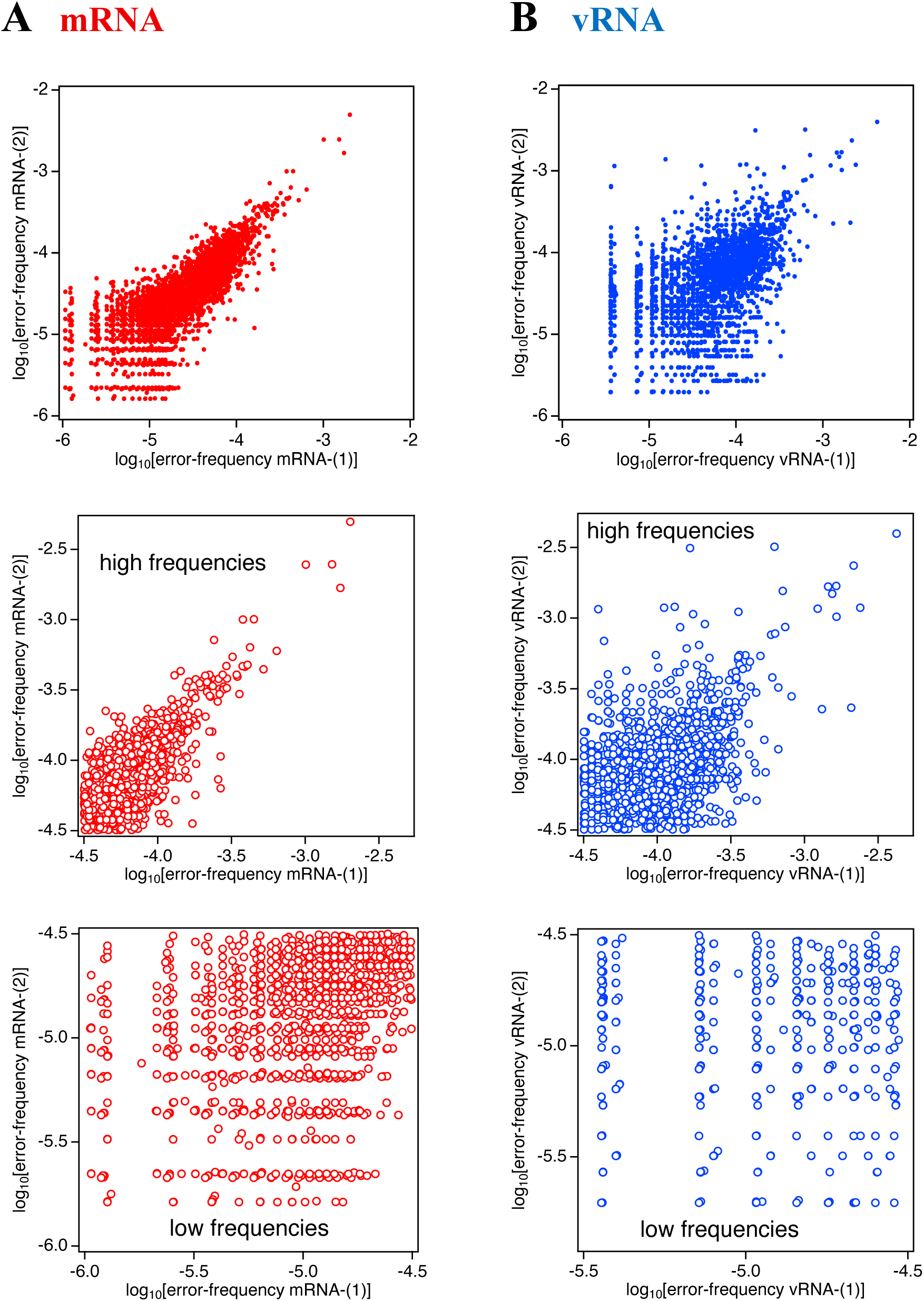
Error landscapes for hemagglutinin-encoding mRNA and vRNA. (A) Decimal logarithm of frequencies for single-site nucleotide changes in hemagglutinin encoding mRNA. Two independent biological samples were studied. Experimental data (Table S3 and S4) are given as plots of logarithm of error frequency for one sample, mRNA-(2), versus logarithm of error frequency for the other sample, mRNA-(1). These plots include the 4360 single-nucleotide changes for which experimental data are available in both replicas. (B) Decimal logarithm of frequencies for single-site nucleotide changes in hemagglutinin encoding vRNA. Two independent biological samples were studied. Experimental data (Tables S5 and S6) are given as plots of logarithm of error frequency for one sample, vRNA-(2), versus logarithm of error frequency for the other sample, vRNA-(1). These plots include the 3060 single-nucleotide changes for which experimental data are available in both replicas. In both, A and B, the plot at the right is the same as the plot at the left, but with axes ranges starting at -4.2, *i.e.,* at a decimal logarithm of frequency roughly corresponding to the error level of the reverse transcriptase used in the sequencing protocol (see text for details).

Briefly (see Methods for details), the virus was prepared through reverse genetics using an eight-plasmid system (Hoffmann et al., 2000) and propagated in MDCK cells. For mRNA sequencing, RNA was extracted from infected cells. We performed sequencing on two totally independent biological samples, mRNA-(1) and mRNA-(2), for which RNA was extracted 2 and 4 hours after infection, respectively. These times were selected in an attempt to favor the mRNA targeted over the vRNA (Phan et al., 2021). In any case, the poly-A- mRNA was purified through poly-A affinity chromatography to eliminate any potential vRNA contamination. The number of UMI families detected per amplicon was about 8-9·10^5^ for the mRNA-(1) sample and about 4-6·10^5^ for the mRNA-(2) sample. The determined error frequencies (Tables S3-S4 and Figure 1A) span several orders of magnitude, from a maximum frequency of about 3·10^-3^ to very low frequencies approaching 10^-6^ (*i.e.*, approaching the value given by ratio of one alternative allele over the number of UMI families). There is a congruence between the data for the two independent biological samples of mRNA (Figure 1A) for frequencies greater than approximately 3·10^-5^, while discrepancies are more apparent at the low frequencies. This pattern will be discussed in detail further below. For a few errors at positions in overlapping amplicon regions, frequencies are determined twice for each of the biological mRNA samples (Table S7). We found an excellent agreement between frequencies determined from different amplicons (Figure S5), which further supports the reliability of our approach.

For vRNA sequencing, we used RNA extracted from virions present in culture supernatants after centrifugation to pellet the virions. Two totally independent biological samples were studied, vRNA-(1) and vRNA-(2). For vRNA-(1), RNA was extracted from the supernatant at 72 hours after infection while, for vRNA-(2), RNA was extracted from the initial supernatant used for infection (formally at time zero). These times were selected in an attempt to favor the vRNA levels (Phan et al., 2021). Affinity chromatography was also used in this case, but not to purify the poly-A-mRNA, but rather to remove any potential mRNA contamination. The results of SafeSeqS sequencing for hemagglutinin-encoding vRNAs are given in Tables S5-S6 and displayed in Figure 1B. Note that mRNA and vRNA have different polarities, but the vRNA results are given for the coding sense. The numbers of UMI families per amplicon identified for vRNA are in most cases substantially smaller (in particular, for amplicon HA3) than the corresponding numbers for mRNA (Table S2). Also, although there is general agreement between the frequency data for the two samples of vRNA studied, the level of congruence appears lower than that observed for the two samples of mRNA, as a visual comparison of Figures 1A and 2A conveys. A lower level of congruence is also apparent when comparing frequencies of errors at overlapping positions in different amplicons (Figure S5).

**Figure 2.**
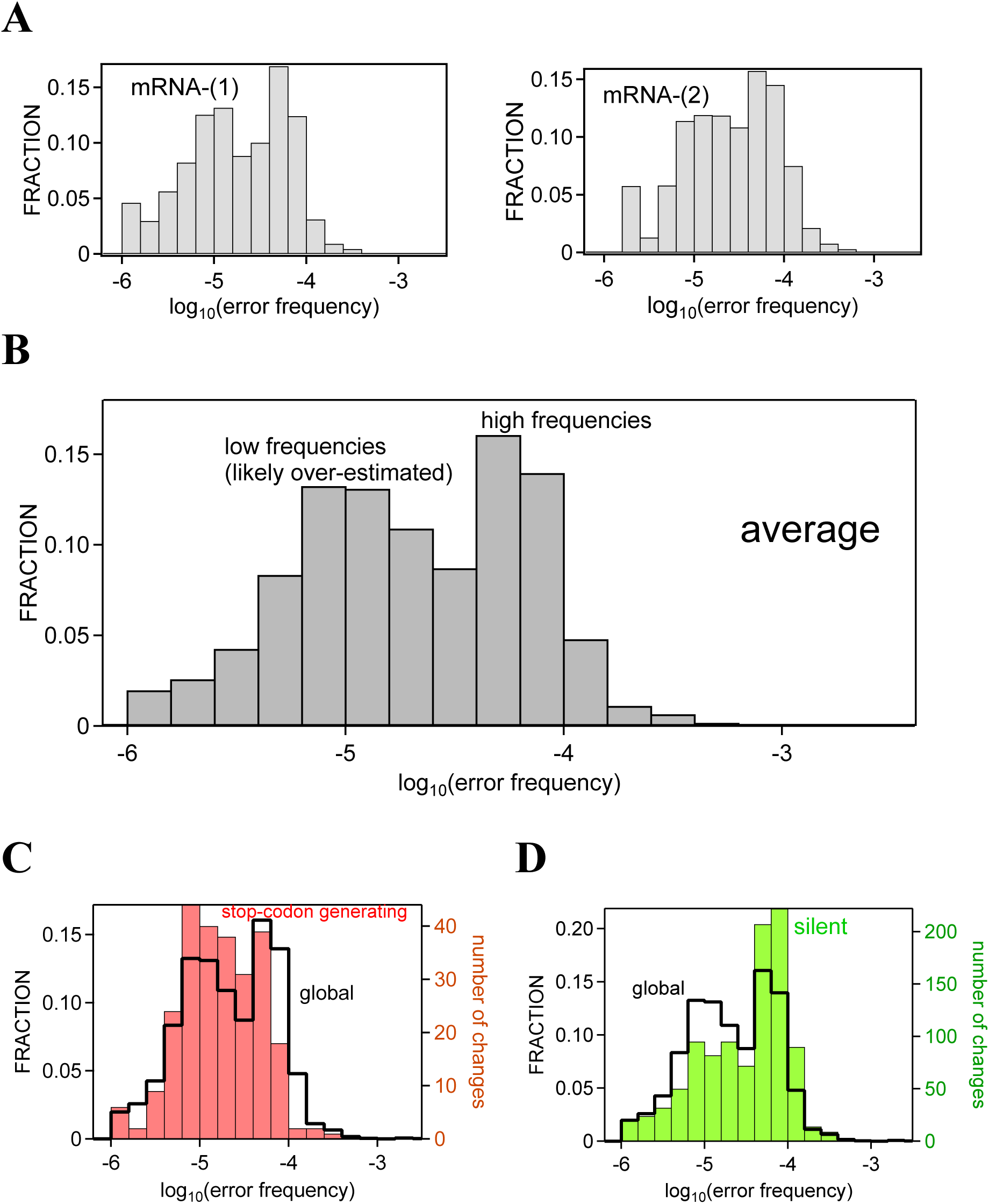
Error frequency distribution for hemagglutinin-encoding mRNA. (A) Distributions for the frequencies of single-site changes derived from the SafeSeqS sequencing of the mRNA-(1) and mRNA-(2) samples. (B) Distribution for the average frequence of the two sets for the mRNA-(1) and mRNA-(2) samples. (C) Distribution derived from the average frequencies, but restricting the calculation to nucleotide changes that generate stop codons (red). (D) Distribution derived from the average frequencies, but restricting the calculation to silent nucleotide changes that do not change the encoded amino acid (green). In B and C the distributions are compared to the global average distribution of panel B, which is shown in cityscape representation. For all distributions, the decimal logarithms of the frequencies are binned, and bin size is 0.2.

Concentration differences may explain, at least in part, the apparent reduced quality of the vRNA sequencing results as compared with mRNA sequencing. RNA concentrations in samples used for mRNA and vRNA sequencing were clearly different: about 1500-2000 ng/μL for the samples used for mRNA sequencing, and about 20-50 ng/μL for the samples used for vRNA sequencing. Even accepting a substantial amount of cellular RNA in the former case, it appears reasonable that the hemagglutinin-encoding RNA abundance is considerably lower in the samples used for vRNA sequencing. The amount of starting material is critical for the first steps of the SafeSeqS protocol, especially for the one-cycle PCR step, where the success of primer-binding and UMI incorporation will be highly dependent on the concentration of target molecules in the mix.

Despite the uncertainties noted above, both the mRNA and vRNA sequencing results are consistent with highly non-random error landscapes. Note also that, as justified in the next section, we will use the mRNA error landscape as a basis for the statistical analyses reported in this work.

### Error frequency distribution for hemagglutinin-encoding RNA

Hemagglutinin-encoding vRNA and mRNA are both synthesized by the viral RNA-dependent RNA-polymerase (Bolvin et al., 2010; Fan et al., 2019). Replication of the vRNA to be encapsulated in the virions involves the synthesis of an RNA copy (cRNA) from the vRNA template followed by the synthesis of a new vRNA from the cRNA template. In this work, we aim at assessing the extent to which influenza evolution/adaptation is determined by non-random diversity in the genetic material. Ideally, the vRNA error landscape should be used for this purpose, because vRNA is inherited and its error landscape should reflect the mutational diversity upon which natural selection acts. However, as noted in the preceding section, our vRNA sequencing data show a much-reduced quality as compared with the mRNA sequencing data. We have, therefore, used the mRNA error landscape as a proxy of the genetic diversity for our analyses. We deem this approach acceptable, since mRNA is synthesized from a vRNA template by the viral RNA-polymerase. That is, the mRNA error landscape should reflect that of vRNA, although additional errors may be introduced during the transcription step. The factors that determine the experimental mRNA error landscape are discussed in the next two sections. Here, we describe the distribution of the error frequencies.

Figure 2A shows the distributions for the frequencies of single-site changes derived from the SafeSeqS sequencing of the mRNA-(1) and mRNA-(2) samples. To obtain these distributions, the decimal logarithms of the frequencies were binned (bin size 0.2) and the number of single-site nucleotide changes with frequencies in each bin were calculated. As was to be expected from the general congruence between the two sets of (Figure 1A), the distributions for the two samples are similar. They both appear bimodal and span frequencies from about 10^-3^ (0.1%) to frequencies about 10^-6^, confirming the error landscape is highly non-random.

Figure 2B shows the distribution for the average frequency of the two sets for the mRNA-(1) and mRNA-(2) samples (see Table S8 for the average values). The distribution for the average values is similar to the individual distributions of Figure 2A. For low frequencies the uncertainties associated to the average values are necessarily large. Yet the average values clearly reflect the pattern of high frequencies vs. low, and likely overestimated (see below), frequencies and provide a useful metric for assessing how the error landscape is sampled in different scenarios.

### Assessing the reverse-transcription distortions in the error-frequency distribution

As noted above (section “SafeSeqS determination of RNA error landscapes”), the UMI-families approach cannot filter out errors generated during the reverse transcription step and during the first amplification cycle in the experimental protocol used. It is, therefore, essential to assess first how such errors, unrelated to RNA biosynthesis, impact our experimental error landscape, as described by the error frequency distribution of Figure 2B.

We used an MMLV-based retrotranscriptase, namely SuperScript IV reverse transcriptase (ThermoFisher Scientific Inc.) with an estimated error rate of around 3.5-6·10^-5^, but even as low as 4.5·10^-6^ for some particular substitutions (ThermoFisher Scientific; Potter et al., 2003; Orton et al., 2015; Jabara et al., 2011; Boutabout et al., 2021;). The DNA-polymerase we used for the DNA amplification step (Phusion High-Fidelity DNA polymerase, ThermoFisher Scientific Inc.) has lower error rate, in the range 10^-7^ (ThermoFisher Scientific; Li et al., 2006) to 10^-6^ (McInerey et al., 2014; Hestand et al., 2016). It seems reasonable, therefore, to assume that the main source of potential distortion in our error landscape comes from the reverse transcription step in the experimental protocol. The low frequency peak in the bimodal distribution of the error frequencies (Figure 2B) appears to be in rough order-of-magnitude agreement with the estimated error rate range for the reverse transcriptase. Plausibly, therefore, distortions produced by the reverse transcription step are mostly confined to the low frequencies, while the high frequency peak in the distribution reflects mostly actual RNA biosynthesis errors (that is, errors generated by the viral RNA-polymerase). This interpretation is consistent with the correlation plots of Figure 1A, which show acceptable congruence between the data for the two mRNA samples for the high frequencies (middle panel), but not for the low frequencies (lower panel).

It is very important to note that the interpretation offered in the preceding paragraph is strongly supported by the analyses reported in this work (see further below), which show that mutations fixed in natural strains over many decades, as well as antibody-escape mutations found in laboratory experiments, preferentially correspond to the high-frequency errors. Such preferential selection would be impossible to rationalize if the high-frequency errors were artifacts generated during reverse transcription. Note also that errors introduced during the reverse transcription step will lead to the low frequencies being overestimates of the actual biosynthesis error frequencies. For instance, a biosynthesis error of negligible frequency may appear with low frequency in our experiments if it is generated during reverse transcription. Overall, the conclusion that the RNA error landscape is highly non-random is robust against reverse-transcription distortions.

### Impact of selection for hemagglutinin function on the error frequency distribution

Once established that the high-frequency errors in our landscape correspond mostly to actual errors generated during RNA biosynthesis, it is important to discuss the factors that determine the high frequency values themselves. Biosynthesis errors are generated by the viral RNA-dependent RNA polymerase, and their frequencies may primarily reflect specific RNA sequence/structure patterns. In addition, however, the frequency values may be modified by selection at the level of vRNA, which is inherited, and the modifications translated to the mRNA which is transcribed from a vRNA template. Under the experimental conditions we used to propagate the virus prior to RNA extraction, we may expect some level of selection related to hemagglutinin function. To explore the contribution from selection to the determined high frequencies, we will consider nonsense and silent changes because they have straightforward functional consequences.

Figures 2C and 2D show distributions derived from the average frequencies of Table S8, but restricting the calculation to nucleotide changes that generate stop codons and to nucleotide changes that do not alter the amino acid encoded. Mutations that generate stop codons preclude the synthesis of functional hemagglutinin and are expected to be subject to strong purifying selection. Mutations that are silent at the amino acid level are expected to be neutral. In both cases (Figures 2C and 2D) the experimental error frequencies span orders of magnitude and define distributions that are similar to the global, average distribution of Figure 2B (shown as a cityscape in Figures 2C and 2D). Yet, Figure 2C indicates that high frequencies are somewhat depleted, but not fully depleted, from stop codons. This pattern supports that stop codons are eliminated through selection but also regenerated through biosynthesis errors. Regarding silent errors, Figure 2D indicates that the high frequencies are somewhat enriched in mutations that do not change the amino acid encoded. This pattern is likely a statistical consequence of the fact that disruptive mutations that impair hemagglutinin function are disfavored, thus leading to enrichment in neutral mutations. Overall, it emerges that the high-frequency peak in the distribution of Figure 2B, reflects mostly errors generated during biosynthesis by the viral RNA polymerase, although the frequency values are modulated to some extent by selection related to hemagglutinin function.

### Error frequency distribution for hemagglutinin at the amino acid sequence level

Figure 2B displays a distribution of error frequencies for single nucleotide changes. However, adaptation is expected to occur at the amino acid sequence level. A one-to-one correspondence between changes at the nucleotide and amino acid sequence levels does not exist, because the genetic code allows many amino acid replacements to be encoded by several (typically 5-7) single nucleotide changes. It is important, therefore, to determine whether the distribution in Figure 2B translates into different amino acid replacement propensities. For instance, if the set of high-frequency, single-nucleotide changes already encoded for most possible amino acid replacements, then the highly non-random nature of the error landscape at the RNA level would probably be of little consequence for adaptation. The simple analysis described below shows this not to be the case.

For each given genetic-code-allowed amino acid replacement in the hemagglutinin of strain A/WSN/1933 H1N1, we have calculated the sum of the error frequencies (Figure 2B and Table S8) for all encoding single-nucleotide changes. This simple calculation provides a useful metric of the probability that the amino acid replacement arises as a consequence of the RNA error landscape. The results of the calculation are given in Table S9 and the corresponding distribution (decimal logarithms binned, bin size 0.2) is shown in Figure 3A. The distribution reproduces the high frequency vs. low-frequency pattern of the RNA error landscape of Figure 2B and the propensities for amino acid replacements span several orders of magnitude. Overall, the distribution shown in Figure 3A can be viewed as describing the consequences of the RNA error landscape at the amino acid sequence level.

**Figure 3.**
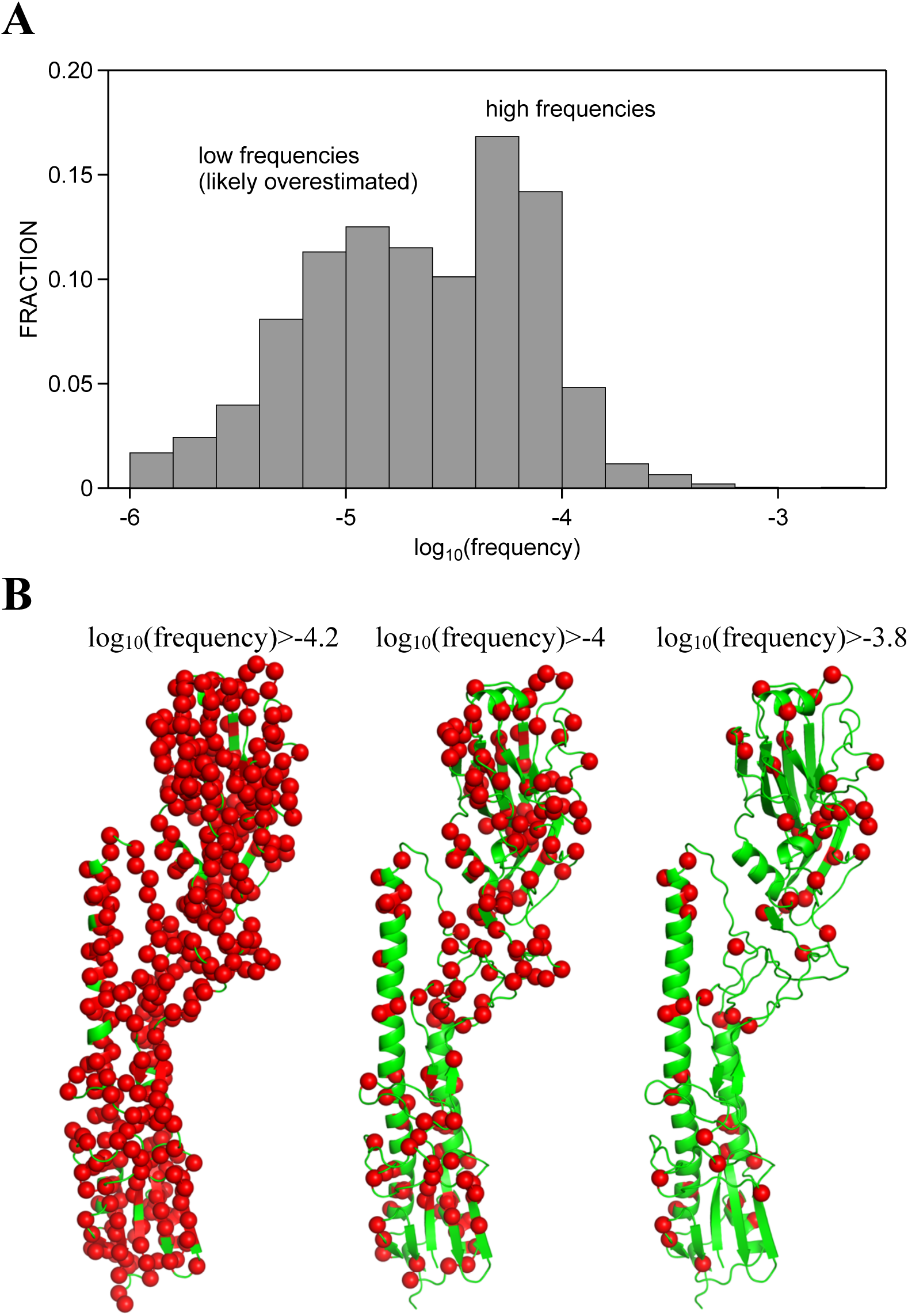
Error frequency distribution for hemagglutinin at the amino acid sequence level. (A) Distribution or error frequencies for amino acid replacements. For each amino acid replacement, the value of an error-frequency metrics is calculated as the sum of the error frequencies for all encoding single-nucleotide changes. The decimal logarithms of the metrics values are binned, and bin size is 0.2.(B) Position-dependence of the propensity for amino acid replacements. The structure of a hemagglutinin monomer is shown in green. To each position in the hemagglutinin amino acid sequence, we assign the highest frequency among all the possible amino acid replacements at the position. Positions with highest frequency values above the threshold shown are highlighted with red spheres.

We have further explored whether the distribution of Figure 3A leads to clearly preferred or to clearly disfavored positions in terms of the probability for harboring amino acid replacements as a result of biosynthesis errors. To this end, we have first assigned to each position in the hemagglutinin amino acid sequence the highest frequency among all the possible amino acid replacements at the position (from Table S9) and then we have highlighted in the hemagglutinin structure the positions with assigned frequencies above given thresholds. We find (Figure 3B) that a pattern of preferred/disfavored positions is apparent only when very high frequency thresholds are used. In fact, with a threshold of log10(frequency)=-4.2, corresponding to the high frequencies in figure 3A, most amino acid sequence positions are susceptible to mutation, although, of course, not all the amino acid replacements will be allowed under that threshold.

The distribution of Figure 3A can be interpreted as describing the consequences of the RNA error landscape at the amino acid sequence level. As such, we will use it in subsequent sections to explore how influenza evolution/adaptation samples the error landscape. The use of the amino acid version of the RNA error landscape will be particularly useful when analyzing antibody escape mutations, because these are often reported in the literature as amino acid replacements (*i.e.,* often, the specific nucleotide changes encoding the evading amino acid replacements are not reported).

### Sampling of the hemagglutinin error landscape during natural influenza evolution

To explore how natural influenza evolution samples the error landscape for hemagglutinin-encoding RNA we have used the following procedure: i) we consider a given natural H1 strain; ii) we align the hemagglutinin amino acid sequence of the natural strain with that of the hemagglutinin from the A/WSN/1933 (see Figure S6 for the alignments); iii) we identify, from the amino-acid-based alignment and the corresponding nucleotide sequences, the nucleotide changes in the hemagglutinin gene for the natural strain with respect to A/WSN /1933 (see Table S10 for the identified nucleotide changes); iv) we assign to each identified nucleotide change the corresponding average frequency in the determined error landscape (*i.e.,* the error frequency given in Table S8) and we calculate the corresponding distribution of frequencies; vi) we compare the calculated distribution for the set of nucleotide changes with the error distribution for all nucleotide changes (Figure 2B). The rationale behind this comparison is straightforward. If the error landscape has no influence in the fixation of mutations during evolution, the two distributions will be similar. On the other hand, if mutations encoded by high-frequency errors are preferentially fixed, the distributions will differ and the high frequencies will be overrepresented in the distribution for the set of nucleotide changes between the two hemagglutinin genes. To make the outcome of the comparison visually intuitive, we have resorted to plots in which both distributions are shown (for instance, Figure 4A) and to violin plots which display smoothed probability densities (for instance, Figure 4B). In fact, we have used the two approaches in all cases, and all the resulting plots are given either in the main text (Figure 4) or in the Supplementary Information (Figure S7).

**Figure 4.**
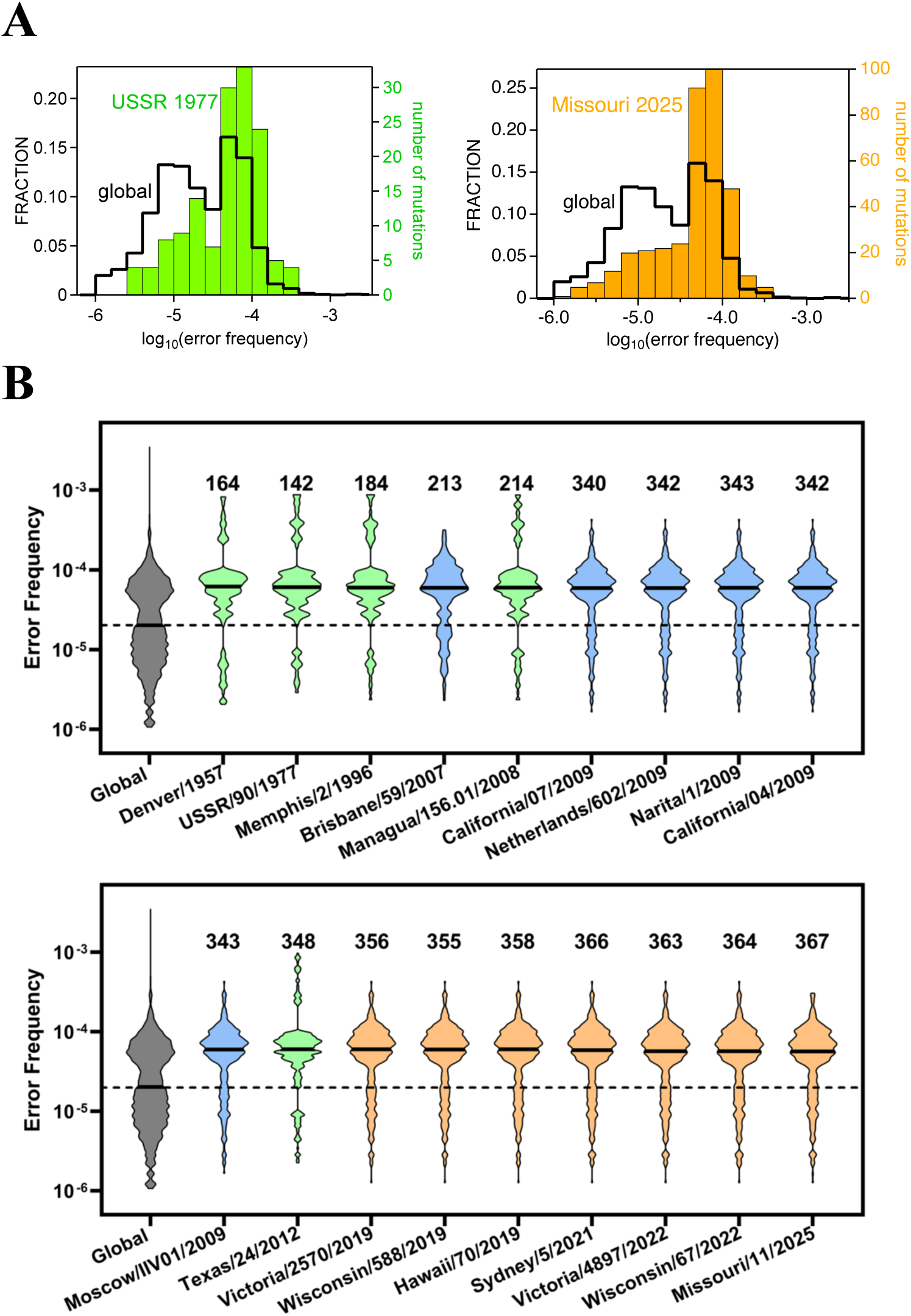
Sampling of the hemagglutinin error landscape during natural influenza evolution. (A) Error frequency distributions for the nucleotide changes in the hemagglutinin gene between natural H1 influenza strains and A/WSN/1933. Decimal logarithms of the frequencies are binned, and bin size is 0.2. The global distribution of Figure 2B is shown here in a cityscape representation. Only two illustrative examples are shown here, but all the calculated distributions are provided in Figure S7. (B) Error frequency distributions for the nucleotide changes in the hemagglutinin gene between natural H1 influenza strains and A/WSN/1933 shown as violin plots. The numbers of nucleotide changes used in each case are shown above the violin. The color code refers to the three strain sets analyzed: green (set-1), orange (set-2) and blue (set-3). The sets are described in the main text, but we note here that set-2 (orange) includes strains recommended by the World Health Organization for inclusion in vaccines for the influenza seasons spanning the period 2019-2025. For comparison the violin plot for the global distribution of Figure 2B is shown in grey. In all cases, the thick horizontal segment represents the median of the distribution. To facilitate global comparison, the median segment for the global (grey) violin has been extended as a dashed line. Violin plots were generated using GraphPad Prism v8.0.1 with the light smoothing option.

We applied the procedure described above to 18 natural influenza strains belonging to three different sets. Set-1) Five strains selected among those included in the phylogenetic analysis of human and swine H1 hemagglutinin sequences provided in Figure 8 of Thyagarajan and Bloom (2014). Set-2) The H1 strains recommended by the World Health Organization for inclusion in vaccines for the influenza seasons spanning the period 2019-2025 (https://www.who.int/teams/global-influenza-programme/vaccines): seven strains. Set- 3) For consistency with the analyses reported in the next section, six strains used in the laboratory experiments on antibody escape.

The numbers of nucleotide changes between hemagglutinin from the selected 18 strains and hemagglutinin from A/WSN/1933 are shown in Figure 4B. For set-1 strains, the numbers roughly increase with the emergence year from about 164 changes (Denver/1957) and 142 changes (USSR/1977) to 348 changes (Texas/2012). On the other hand, most set-2 and set-3 strains show about 340-360 changes in the hemagglutinin gene with respect to A/WSN/1933. Actually, the hemagglutinins for the set-3 strains (except for Brisbane/2007) display high sequence identity with each other and appear closely related (see Figure S8) to the hemagglutinin of the pandemic A/H1N1pdm09 viruses that emerged in 2009 in the United States and Mexico (Smith et al., 2009). Furthermore, the hemagglutinins for set-2 strains (strains recommended by the World Health Organization for inclusion in vaccines for the influenza seasons spanning the period 2019-2025) appear to have evolved from pdm09 hemagglutinin (see Figure S8). Beyond these specific sequence details, however, our analyses (Figures 4 and S7) reveal preferential fixation of mutations encoded by the high-frequency errors in all cases. That is, the fraction of high frequencies is consistently increased in the subset of errors encoding for fixed mutations in the hemagglutinin gene (Figures 4 and S7).

The clear message conveyed by the visual inspection of Figures 4 and S7 is confirmed by Kolmogorov-Smirnov tests. Values of the KS statistics (the maximum absolute difference between cumulative distributions) can be easily calculated by comparing the distributions for the nucleotide changes fixed in given strains with the global distribution for all nucleotide changes. The values thus calculated are within the range 0.28-0.35. The number of nucleotide changes vary between 142 and 367 (Figure 4B), which leads to critical values of the KS statistics within the 0.10-0.16 range for a confidence level of α=0.001 (https://real-statistics.com/statistics-tables/kolmogorov-smirnov-table/). The experimental KS statistics (*i.e.,* the values derived from the pairwise comparison of distributions) are thus larger than the critical values for α=0.001. Therefore, the null hypothesis that the fixed changes in the hemagglutinin gene reflect random sampling of the global distribution can be rejected for all natural strains tested at confidence levels even lower than 0.001.

The analyses we have carried out (Figures 4 and S7) are obviously subject to several uncertainties and problems that could potentially blur the statistical signal of preferential selection. First, they implicitly use the hemagglutinin from the A/WSN/1933 strain (or the closely related A/Wilson-Smith/1933 strain) as the ancestor of the hemagglutinin from the natural strains under consideration. This may be a good approximation in some cases, but perhaps not in other cases. For instance, according to the phylogenetic tree of human and swine H1 hemagglutinin sequences provided in Figure 8 in Thyagarajan and Bloom (2014), hemagglutinin from A/Managua/156.01/2008 shares a common ancestor with the hemagglutinin from the A/Wilson-Smith/1933, and this common ancestor is very close to the later. On the other hand, hemagglutinin from A/Texas/24/2012 shares a common ancestor with hemagglutinin from A/Wilson-Smith/1933, but this common ancestor, related to the 1918 flu, is far more distant. Clearly, differences between the sequence of hemagglutinin from A/WSN/1933 strain (or the closely related A/Wilson-Smith/1933) and the relevant ancestor will introduce spurious nucleotide changes in the analysis. Second, some of the identified nucleotide changes may have resulted from “double hits”, that is, from two changes at the same position occurring along the evolutionary trajectory, with the concomitant distortion of the statistical analysis. Third, our calculations implicitly assume that the error landscape determined for hemagglutinin-encoding RNA from A/WSN/1933, as determined under the specific conditions of our experiments and analyses, applies to hemagglutinin from strains that emerged long after 1933.

It is remarkable that, despite the several issues noted in the preceding paragraph, we consistently find a clear statistical preference when considering natural strains spanning many decades. This outcome supports that the preferential selection of mutations encoded by high-frequency errors is a robust feature, and that the error landscape is evolutionarily conserved to some substantial extent. Plausibly, RNA sequence/structure patterns responsible for error promotion are not drastically altered upon the accumulation of a moderate number of nucleotide changes.

Finally, we have carried out an analysis similar to that reported above (Figures 4 and S7) but at the amino acid sequence level. We thus identified from the amino acid sequence alignments (Figure S6), the hemagglutinin sequence differences between each of the 18 strains and A/WSN/1933, calculated the corresponding distributions using the error metric for amino acid replacements of Table S9 and compared these distributions with the global distribution of Figure 3A. This amino-acid-level analysis is subject to additional uncertainty because we do not consider the specific errors encoding the amino acid replacements but, rather, a general metric of the probability that the amino acid replacement arises as a consequence of the RNA error landscape. Still, the analysis (see Figure S9) does support preferential fixation of amino acid replacements encoded by high-frequency errors.

### Sampling of the hemagglutinin error landscape by antibody escape mutations

Laboratory experiments on antibody escape confront a natural virus, or an engineered viral population in the case of deep mutational scanning (DMS) experiments, to neutralizing antibodies and determine the mutations at the amino acid sequence level that enable virus propagation. In DMS experiments, the mutational landscape sampled by the virus is determined largely by the engineered library of viral variants. Therefore, we will be concerned in this section with laboratory experiments on antibody escape with natural virus strains, although, for comparison, we will also discuss a DMS experiment. The specific experiments we analyze are summarized in Table S11 in terms of the virus strain, antibodies and number of escape mutations identified. The studies collected in Table S11 were derived from an extensive literature search, as described in Methods. Interestingly, except for Brisbane/2007, all these studies (see Figure S8) used strains related to the pandemic A/H1N1pdm09 viruses that emerged in 2009 in the United States and Mexico (Smith et al., 2009). Also, all these studies reported the antibody escape mutations only as amino acid replacements (*i.e.,* the encoding nucleotide changes are not available). Therefore, our analysis must use the general metric of the probability that the amino acid replacement arises as a consequence of the RNA error landscape (Figure 3A and Table S9), instead of the original error frequencies for nucleotide replacements (Figure 2B and Table S8). Despite this additional source of uncertainty, the statistical signal of preferential selection is clearly detected.

Among the studies described in Table S11, Matzusaki et al. (2014) reported 43 escape mutations, 23 of which could be used for our analysis. All other articles in Table S11 (O’Donell et al., 2012; Rudneva et al., 2012; Schmeisser et al., 2013; Retamal et al, 2014; Yasuhara et al., 2017; Prachanronarong et al., 2019; Gao et al., 2020; Lee et al., 2023) report just a few escape mutations each. Therefore, for the purpose of our statistical analyses, we constructed two sets of mutations. One set included solely the escape mutations from Matzusaki et al. (2014), while an expanded included, in addition, the escape mutations from all other articles in Table S11. To explore how the hemagglutinin error landscape is sampled by the antibody escape mutations, we performed an analysis similar to that described in the preceding section for the natural strains. That is, we calculated the distribution of error frequencies for the two sets of antibody-escape amino acid replacements and compared the calculated distributions with the distribution for all the possible amino acid replacements that is shown in Figure 3A. The comparison (Figure 5A) indicates that most antibody-escape mutations correspond to high-frequency errors in the landscape. This result is also apparent from the violin plots of Figure 5B. Furthermore, according to Kolmogorov-Smirnov tests, the null hypothesis that the sets of antibody escape mutations reflect random sampling of the global distribution is rejected at confidence level of α=0.05 for the set based on Matzusaky et al. (2014) and at confidence level of α=0.01 for the expanded set.

**Figure 5.**
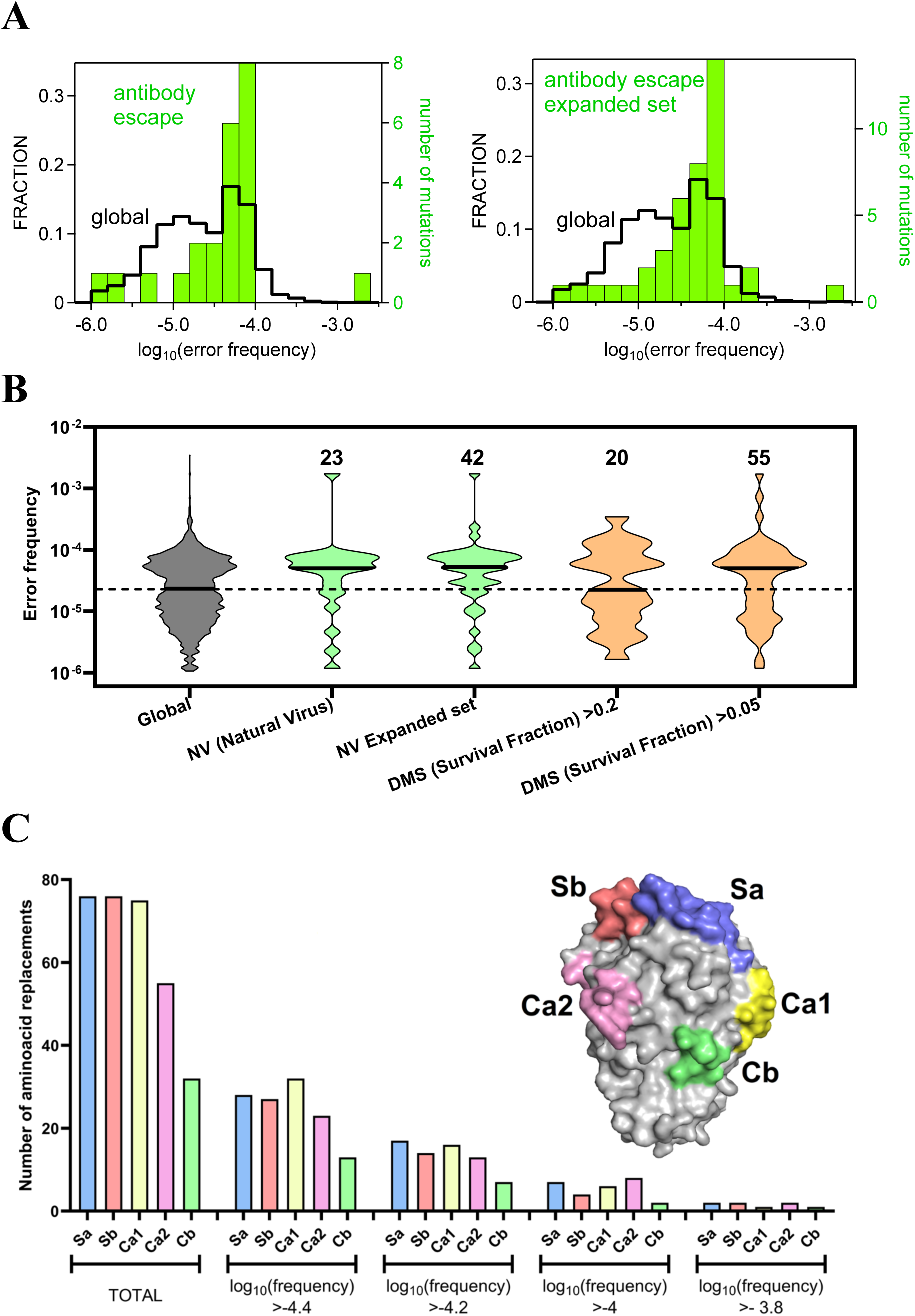
Sampling of the hemagglutinin error landscape by antibody escape mutations. (A) Error frequency distributions for escape mutations determined in laboratory experiments on antibody evasion with neutral influenza strains (Table S11). The distribution of the plot at the left is calculated for 23 escape mutations reported by Matzusaki et al. (2014), while the expanded set used for the distribution in the plot at the right includes, in addition, 21 escape mutations reported in various articles in the literature (see text for details). Decimal logarithms of the frequencies are binned, and bin size is 0.2. For comparison, the global distribution of Figure 3A is shown here in a cityscape representation. (B) Error frequency distributions for sets of antibody escape mutations shown as violin plots. The number of amino acid differences used for each strain is shown as a number above the violin. The distributions of mutations derived from laboratory experiments with natural strains are also shown here in green. In addition, we show (orange) distributions calculated for mutations determined in deep mutational scanning experiments (DMS; Doud et al., 2018) for two different survival fractions (Table S12). For comparison the violin plot for the global distribution of Figure 3A is shown in grey. In all cases, the thick horizontal segment represents the median of the distribution. To facilitate global comparison, the median segment for the global (grey) violin has been extended as a dashed line. Violin plots were generated using GraphPad Prism v8.0.1 with the light smoothing option. (C) Number of amino acid replacements at positions corresponding to antigenic sites in the hemagglutinin head under different error-frequency thresholds. TOTAL refers to the total number of amino acid replacements allowed by the genetic code for single-nucleotide changes. The calculation is carried out for the antigenic sites reported by Dzimianski et al. (2023), which are highlighted in the structure shown for the hemagglutinin head.

For comparison, we have also analyzed the results of deep mutational scanning (DMS) experiments on hemagglutinin from the A/WSN/1933 strain (Doud et al., 2017; Doud et al. 2018). Very briefly, virus libraries were constructed using a protocol that generates a few random codon mutations per gene (Bloom, 2014), were then exposed to antibodies and, twelve hours after transfer to growth medium, deep sequenced. Results were reported (Doud et al., 2018) as viral survival fractions associated to different amino acid mutations. The distributions of the error frequencies from Table S9 for mutations that were reported to achieve survival fraction ≥0.2 and survival fraction ≥0.05 (see Table S12) are given in Figure 5B. For survival fraction ≥0.2 the distribution is congruent with the global distribution for all amino acid replacements of Figure 3A, plausibly reflecting that the critical antibody neutralization step samples the initial random variant distribution. On the other hand, under the more relaxed selection involving survival fraction ≥0.05, the distribution reveals again preferential sampling of the low-frequency mutations in the error landscape. Plausibly, this result reflects the sampling of the actual hemagglutinin error landscape during the growth period in the experimental protocol.

The analyses reported in this section show that antibody-evading mutations emerging in natural influenza viruses, but not necessarily the mutations found in DMS experiments on engineered viral libraries, preferentially correspond to high-frequency changes in the error landscape of hemagglutinin-encoding RNA. It follows that, together with factors such as the requirement to maintain viral fitness and the capability to disrupt antibody binding (Thadani et al., 2023), the likelihood that a mutation is fixed in response to prior infection or vaccination depends on the frequency of the encoding error. Note that virus propagation can indeed be enabled by errors at comparatively low level (Luzon-Hidalgo et al., 2025).

It follows that approaches to predict evasion and influenza evolution in general could benefit from the information provided by biosynthesis error landscapes. This is the more so when considering that the error frequencies span orders of magnitude and, consequently, that they can be used to establish a clear a priori ranking of mutation propensities. To illustrate this point in a simple manner, we show in Figure 5C the number of amino acid replacements at positions corresponding to antigenic sites in the hemagglutinin head (Dzimianski et al., 2023) under different error-frequency thresholds. It is apparent that even low error-frequency thresholds drastically reduce the number of amino acid mutations with respect to the total allowed by the genetic code for single-nucleotide changes.

## CONCLUDING REMARKS

Natural selection acts on diversity generated through errors in the biosynthesis of the genetic material. Two crucial questions thus arise: To what extent is the biosynthesis error landscape non-random? Does a non-random error landscape determine to some extent future evolution and adaptation? Here, we have addressed these two questions in the context of influenza evolution/adaptation linked to changes in the hemagglutinin molecule.

Our experimental results show that the error landscape for hemagglutinin-encoding RNA is highly non-random, with frequencies for individual errors spanning orders of magnitude. This wide frequency range plausibly reflects that errors are promoted/depleted by specific RNA sequence/structure patterns.

We find that antibody-escape mutations from laboratory experiments preferentially correspond to high-frequency errors. We obtain the same general result upon analysis of hemagglutinin changes in natural influenza strains over several decades. Preferential sampling of mutations encoded by high-frequency errors is plausibly a probabilistic effect. Assume, for instance, that two different amino acid replacements evade a given antibody and that they are encoded by errors with quite different frequencies. The replacement encoded by the higher-frequency error is more likely to be fixed in the presence of the antibody, because it is represented at a higher level in the initial viral population.

There is considerable interest in developing approaches to forecast influenza virus evolution. Our results and analyses support that, together with the capability to disrupt antibody binding while maintaining viral fitness, the likelihood that a mutation is fixed in response to immune reaction will depend on the frequency of its encoding error. It follows that computational approaches to anticipate virus evolution should benefit from the information provided by biosynthesis error landscapes.

Recurring influenza infections over successive years and the low effectiveness of influenza vaccines are generally put down to the rapid evolution of influenza viruses. Our results and analyses reveal a factor potentially contributing to such rapid evolution. Namely, the fact that the error landscape for hemagglutinin-encoding RNA is highly non-random and includes a substantial fraction of high-frequency errors. Antibody-evading mutations encoded by high-frequency errors will display enhanced fixation probability thus promoting fast adaptation to the host immune response. The existence of this mechanism could have an impact in vaccine design.

## Data availability

All calculations reported in this paper can be reproduced from the raw sequencing data deposited in Zenodo under DOI: 10.5281/zenodo.20622529, together with the methodological details and supplementary tables provided with the article. The Zenodo record includes the raw paired-end FASTQ files, the Sanger sequencing files used to identify primer-binding regions, sample metadata, checksums, and a README file describing the dataset structure.

## Acknowledgments

This work was supported by Grant IHRC22/00004 (to J.M.S.-R.) funded by the “Instituto de Salud Carlos III” and Next Generation EU.

## METHODS

### Cell Culture

HEK293T/17 human embryonic kidney cells and Madin–Darby canine kidney (MDCK) cells were supplied by the Cell Bank of the Centro de Instrumentación Científica (CIC) at the University of Granada (Granada, Spain). Cells were maintained in Dulbecco’s Modified Eagle Medium (DMEM; Gibco) supplemented with 10% fetal bovine serum (FBS; Gibco), 25 mM HEPES (Gibco), 100 U/mL penicillin (Sigma-Aldrich), and 100 μg/mL streptomycin (Sigma-Aldrich). Cell cultures were incubated at 37°C in a humidified atmosphere containing 5% CO₂ and routinely passaged to maintain exponential growth.

### Cell Transfection and Viral Supernatant Collection

Reverse genetics rescue of influenza A virus was performed using eight plasmids derived from the A/WSN/1933 (H1N1) strain (Hoffmann et al., 2000), provided by the St. Jude Children’s Research Hospital (Memphis, Tennessee): pHW181-PB2, pHW182-PB1, pHW183-PA, pHW184-HA (HA: hemagglutinin), pHW185-NP, pHW186-NA, pHW187-M and pHW188-NS. For virus rescue, HEK293T/17 and MDCK cells were co-cultured in six-well tissue culture plates (Thermo Fisher Scientific). Cells were seeded at a density of 5 × 10⁵ cells per well of each cell type (total of 1 × 10⁶ cells per well) in DMEM supplemented with 10% FBS, 25 mM HEPES, penicillin (100 U/mL), and streptomycin (100 μg/mL). Co-cultured cells were transfected with the eight plasmids using Lipofectamine 3000 (Invitrogen) according to the manufacturer’s instructions. Briefly, 312.5 ng of each plasmid was used per well, and transfection complexes were prepared in Opti-MEM reduced-serum medium (Gibco). As transfection controls, three additional experimental situations were established: i) Without pHW182-PB1 plasmid (S/PB1); ii) Without pHW184-HA plasmid (S/HA); iii) Adding PBS in the plasmid’s place. As described in the following section, cell death did not occur in the transfection controls. Cells were incubated with the transfection mixture for 4.5 hours at 37°C under 5% CO₂. After incubation, cells were washed twice with phosphate-buffered saline (PBS; Gibco) to remove residual transfection reagents, and the medium was replaced with DMEM supplemented with 5% FBS and 25 mM HEPES, without antibiotics. At 24 hours post-transfection, cell culture supernatants were collected and stored at 4°C, and fresh medium was added to the cells. A second collection was performed at 48 hours post-transfection, and both supernatants were pooled. Collected supernatants were clarified by centrifugation at 300 × g for 5 minutes at 4°C and subsequently filtered through 0.22 μm pore-size filters. Viral and control supernatants were aliquoted and stored at −80°C until further use.

### Infection and Viral mRNA Extraction

MDCK cells were seeded in 75 cm² flasks (Thermo Fisher Scientific) at a density of 5 × 10⁶ cells per flask and incubated for 24 hours prior to infection. Cells were then infected with viral supernatants at a titer of 2.2 × 10⁸ TCID₅₀/ml and incubated for 1 hour at 37°C under 5% CO₂ to allow viral adsorption. Additionally, cells were mock-infected with the supernatants for the controls described above. Following infection, viral inoculum was removed, and cells were washed with PBS and incubated in DMEM supplemented with 5% FBS and 25 mM HEPES, without antibiotics. At 2, 4 and 6 hours post-infection, cell culture supernatants were removed, and total RNA was extracted from MDCK cells using the RNeasy Mini Kit (Qiagen), following the manufacturer’s instructions. These times were selected in an attempt to favor the mRNA targeted over the vRNA (Phan et al., 2021). Antigen-tests for influenza A virus were done to supernatants of infected and mock-infected cells, resulting in positive and negative results, respectively (Figure S10). In addition, optical microscope images of cultures were taken (Figure S11) and showed cell death with the sample of virus-infected cells and alive adherent cells with the mock controls. Finally, the purified RNA samples were sent in dry ice to AllGenetics & Biology SL (Spain), for further processing and sequencing analysis where they were immediately transferred to an ultrafreezer (-80 °C) to preserve RNA integrity.

### Viral RNA (vRNA) extraction from viral particles

To obtain viral RNA (vRNA) from viral particles, an additional infection was performed using MDCK cells. MDCK cells were cultured and subsequently infected with viral supernatants as previously described. After 72 hours of incubation (vRNA-(1)), cell culture supernatants (SN) were collected. In parallel, a supernatant derived from the transfection step was included as a 0-hour infection control (vRNA-(2)). Both supernatants were clarified by centrifugation at 17,000 × g for 10 minutes at 4°C to remove cell debris, followed by filtration through 0.22 μm pore-size filters. Clarified and filtered supernatants were then subjected to ultracentrifugation at 120,000 × g for 4 hours at 4°C to pellet viral particles. The resulting pellet was resuspended, and viral RNA was extracted using the RNeasy Mini Kit (Qiagen), following the manufacturer’s instructions. Subsequent steps of RNA processing and analysis were performed as described above.

### SafeSeqS library preparation and sequencing

In order to sequence the viral hemagglutinin transcripts, we designed a set of four primer pairs that span the whole hemagglutinin genetic sequence in fragments of around 400-500 bp each. The genetic sequence upon which the primers were designed was determined by Sanger sequencing. Reverse transcription PCR (RT-PCR) was performed to detect the hemagglutinin (HA) gene using the SuperScript IV One-Step RT-PCR System (Invitrogen) under the following conditions: a reverse transcription step at 50°C for 10 minutes, initial denaturation at 98°C for 5 minutes, followed by 39 cycles of 98°C for 10 seconds, 52°C for 10 seconds, and 72°C for 15 seconds, and a final extension at 72°C for 5 minutes. PCR product size was verified by agarose gel electrophoresis (Figure S12). The primers used for amplification were as follows: forward primer, 5′-GCAGGGGAAAATAAAAACAACC-3′, and reverse primer, 5′-TGGAAGCAGTGAGTCGCAT-3.

Primers were designed to ensure compatibility among primer pairs, avoid the formation of secondary structures, and provide overlap between consecutive amplicons. Then, a 12-bp random sequence (UMI) and Illumina sequencing primer sequences were added to the 5′ end of each primer (Table S1).

For mRNA sequencing, samples were enriched in Poly-A mRNA in order to separate any encapsulated viral genome present from the transcripts. For this, we used the mRNA Magnetic Isolation Module (NewEngland Biolabs), using 1 μL of the total RNA as input. We strictly followed the manufacturer’s instructions up until the final elution step, which was slightly modified to make the product compatible with the downstream process.

For vRNA sequencing, the protocol of the NEBNext Poly(A) mRNA MagneticIsolation Module (New England Biolabs) was applied to 5 μL of the sample. However, in this case, any poly-A mRNA present would bind to the Oligo d(T)25 beads, which was discarded, and the remaining vRNA in the supernatant was retained. This fraction was subsequently purified using the NucleoMag NGS Clean-up and size selection (Macherey-Nagel).

For library preparation, we retrotranscribed Poly A-enriched RNA and viral RNA samples in four separate reactions each, using the reverse primer from each pair described above, and the SuperScript IV Reverse Transcriptase (Thermo Fisher Scientific), under the experimental conditions indicated by the manufacturer. The reaction was stopped after 10 minutes by incubating the samples at 80 °C for 10 minutes. After cDNA synthesis, leftover RNA was removed by incubating the samples at 37 °C for 20 minutes in the presence of RNase H. The resulting cDNA was amplified by performing a 1-cycle PCR in order to incorporate the forward UMI and sequencing primer.

The PCRs to generate the four amplicons from the mRNA were carried out in a final volume of 12.5 μL, containing 5.95 μL of template cDNA, 0.48μM of the forward primer, and 6.25 μL of Phusion High-Fidelity PCR Master Mix (ThermoFisher Scientific). The reaction mixture was incubated as follows: an initial denaturation step at 98 °C for 2 min, followed by 1 cycle of incubation at the specific annealing temperature for each primer (A1: 59 °C, A2: 62 °C, A3: 65 °C, A4: 58.6 °C) for 2 min, and a final extension step at 72 °C for 10 min. The PCR products were then purified using the MagBind RXNPure magnetic beads (Omega Biotek), following the manufacturer’s instructions.

The PCRs to generate the four amplicons from the vRNA (Table S1) were carried out in a final volume of 12.5 μL, containing 2 μL of template cDNA, 0.48μM of the forward primer, and 6.25 μL of Phusion High-Fidelity PCR Master Mix (ThermoFisher Scientific), and ultrapure water up to 12.5 μL. The cycling conditions were identical to those described above for the mRNA-derived amplicons. The PCR products were then purified using the MagBind RXNPure magnetic beads (Omega Biotek), following the manufacturer’s instructions.

Uniquely UMI-tagged molecules were then amplified in a second PCR round in order to make multiple copies for each UMI combination (UMI families). This second PCR was carried out in a final volume of 25 μL, containing 5 μL of template cDNA (the product from the first PCR), 0.2 μM of both the forward and reverse primers, 13.75 μL of Q5® High-Fidelity 2X Master Mix (New England Biolabs), and ultrapure water up to 25 μL. The reaction mixture was incubated as follows: an initial denaturation step at 98 °C for 30 s, followed by 30 cycles of 98 °C for 10 s, 60 °C for 30 s, and 72 °C for 30 s; followed by a final extension step at 72 °C for 7 min.

In this second PCR, samples were also dual-indexed so they could be pooled together for sequencing and demultiplexed after sequencing. PCR products were purified twice following the magnetic bead protocol, as described above. A negative control that contained no DNA was included in every PCR round to check for contamination during library preparation. Purified libraries were pooled in equimolar amounts according to the results of a Qubit dsDNA HS Assay (Thermo Fisher Scientific) quantification. The pool was sequenced in a fraction of a NovaSeq PE250 flow cell (Illumina) aiming for a total output of 24 gigabases.

### Quality control and pre-processing of the raw sequencing data

Illumina paired-end raw data for each library consists of forward (R1) and reverse (R2) reads stored in separate files, which also include the reads’ quality scores. We assessed the quality of the FASTQ files with the software FastQC v0.11.9 (Andrews, 2010) and summarised the output using MultiQC v1.19 (Ewels et al., 2016).

The pre-processing of the raw data was performed as follows. First, Trimmomatic v0.39 (Bolger et al., 2014) was used to remove sequences containing adapters (ILLUMINACLIP option followed by a requirement for a minimum length of 250), low quality regions (AVGQUAL:28 and SLIDINGWINDOW:15:30) and sequences shorter than 180 base pairs after this quality filtering step. Second, Cutadapt v4.2 (Martin, 2011) was used to remove reads that did not contain the primer sequences (allowing up to 10% mismatches relative to primer length). Then, the fastp software v0.23.3 (Chen et al., 2018) was used to perform a read correction step, during which R1 and R2 reads are overlapped, and, in case of any mismatched base pair, the base with the highest quality score is assigned to both reads. Finally, the MEM algorithm (Li, 2013) of BWA v0.7.17-r1198-dirty (Li and Durbin, 2009, 2010) was used to map quality-trimmed reads to the hemagglutinin gene. Then, selection of unique, non-long soft-clipped reads was achieved using SAMtools v.1.19 (Li et al., 2009) and grep command of SeqKit toolkit v2.6.1 (Shen et al., 2016). This process ensured the exclusion of low-quality alignments in subsequent analysis.

The quality of the filtered reads was checked again using FastQC and MultiQC to make sure that only high-quality reads were used in the subsequent analysis.

### UMI detection and family assembly

UMI detection and family assembly were performed using MAGERI v1.1.1 (Shugay et al., 2017), an all-in-one pipeline specifically designed for processing and genotyping UMI-tagged data. MAGERI was run using -M2 demultiplexing module and default presets, with the exception of two parameters that were modified: defaultOverseq was changed from 5 to 2 (it specifies the minimum number of reads supporting a UMI family for inclusion in the subsequent analysis) and readLength was adjusted from 100 to 250. Also, a metadata file containing forward and reverse primer sequences used for recovering each amplicon was required.

An overview of the pipeline is as follows. First, MAGERI performs UMI extraction and demultiplexing. UMI sequences are extracted from the reads, discarding those with a minimal quality below a specified threshold (Phred 20) and reads tagged with identical UMI sequences are grouped into molecular identifier groups (MIGs) or families. Only UMI families with the minimum support criteria (≥2) are kept. Subsequently, demultiplexed reads are used as input for the assembly of a consensus sequence for each UMI family. Bases must have quality values greater than 30 and frequency values over 10^-3^ to be considered when calculating the consensus quality score. The consensus sequence generation aims to remove errors introduced during sequencing. Assembled consensus sequences were then mapped to the hemagglutinin gene of the Influenza A virus (strain A/WSN/1933(H1N1)) which had been obtained through Sanger sequencing. Although MAGERI also performs variant calling, an independent variant calling step was carried out using MAGERI-generated mappings to provide greater control over the analysis and variant call outputs.

### Variant calling

We performed variant calling using the MAGERI-generated mappings. First, overlapping regions between paired-end reads were clipped using clipOverlap option from bamUtil v1.0.15 toolkit (Jun et al., 2015), ensuring that bases coming from the same initial molecule were not double-counted when estimating variant allele frequencies. Then, as an additional quality-control step, mappings were filtered to remove secondary, unmapped and QC-failed alignments before BAM sorting and indexing with SAMtools.

Variant detection was then conducted with BCFtools v1.19 (Danecek et al., 2021). First, we used the mpileup command to generate genotype likelihoods at each genomic position with coverage, applying a minimum base-quality threshold of 38 (--min-BQ 38), interpreting MAGERI’s Consensus Quality Score (CQS) as base quality. The threshold of CQS ≥ 38 retains only bases supported by ≥ 96.25% of reads within each UMI family, thereby implementing the SafeSeqS concordance criterion (Kinde et al., 2011), but with a slightly more stringent cutoff. Also, probabilistic realignment was disabled (-B) to prevent it from modifying base quality values representing within-family concordance.

Variants were then genotyped with the BCFtools call module, disabling indel calling, retaining all alternate alleles, and applying the multiallelic rare-variant caller (--skip-variants indels --keep-alts --multiallelic-caller options). No additional post-calling filters were applied beyond the CQS threshold. For each sample and amplicon, multiallelic records were decomposed to one line per alternate allele prior to annotation. The final set of SNVs was then annotated using SnpEff5.1f (Cingolani et al., 2012), a software toll for variant effect predictor. SnpEff reports details on the functional class of analyzed variants, their gene impact, codon and amino acid changes, with additional annotations. Annotation required a reference genome database, which was generated from the ‘LC333185.1’ NCBI entry.

### Literature search for antibody-escape mutations

We performed a literature search using Google Scholar and PubMed with the following keywords “H1N1 influenza escape mutants”, “antibody escape mutants influenza H1N1”, “Hemagglutinin H1 escape mutants”, “H1N1 escape mutants viral strains” and “Neutralization assays H1 escape mutants”. This search led to about 100 articles, published, in most cases, between 2019 and 2026. However, we excluded most of these articles because they did not directly address antibody escape or did not report clearly identifiable mutations on H1 strains. Actually, only one of the articles, Matzusaki et al. (2014), fulfilled our selection criteria. For this reason, we performed a secondary search on the articles that appear as “citing” or as “related to” Matzusaki et al. (2014) in Google Scholar and PubMed. We thus examined about 100 additional articles. While the majority were discarded for the same reasons mentioned above, 8 did fulfill our selection criteria. Overall, we found 9 articles that reported 98 different antibody-escape mutations (Table S11). Not all these mutations were useful for our analyses, however, because often the background amino acid residue in the hemagglutinin of the strain studied did not match the corresponding residue (*i.e.,* the residue at the same position after alignment) in our reference hemagglutinin from A/WSN/1933. Also, some mutations were reported in two or more studies using different strains. Overall, the analyses described in the main text involve a total of 42 antibody-escape mutations.

### Structural modeling

For the structure of the hemagglutinin from the A/WSN/1933 strain (UniProt:A0A2Z5U3Z0), we used a SWISS-MODEL model (ID:E6XTV3) based on the crystallographic structure 6N41 which corresponds to the hemagglutinin of the A/Netherlands/002P1/1951 (sequence identity of 88% with hemagglutinin from A/WSN/1933). Images of 3D-structures (Figures 3B and 5C) were generated with PyMOL Molecular Graphics System, Version 3.1.6.1 (Schrödinger, LLC).

