## Supporting Information for "Influenza evolution/adaptation samples a highly non-random error landscape for hemagglutinin-encoding RNA"

[1]Departamento de Quimica Fisica. Facultad de Ciencias, Unidad de Excelencia de Quimica Aplicada a Biomedicina y Medioambiente (UEQ), Universidad de Granada, 18071 Granada, Spain.
[2] AllGenetics & Biology SL, Calle Cubelos 21, Oleiros, 15173 A Coruña, Spain.
[3]Departamento de Parasitología, Facultad de Ciencias, Universidad de Granada, 18071 Granada, Spain.

**Table S1. Primers used in hemagglutinin SafeSeqS sequencing**

| AMPLICON | Primer name | F/R | |
|---|---|---|---|
| HA1<br>26-431 | HA1_35F_corr | Forward | GAAGGCAAAACTACTGGTC |
| | HA1_492R | Reverse | GGGAGCATGATACTGTTAC |
| HA2<br>431-847 | HA2_440F | Forward | GGAAAGTTCATGGCCCAAC |
| | HA2_910R | Reverse | GTGTTACACTCATGCATTGAC |
| HA3<br>848-1268 | HA3_857F | Forward | AGGGTTTGAGTCCGGCATC |
| | HA3_1331R | Reverse | GTCCAGAAACCCATCATCAAC |
| HA4<br>1247-1650 | HA4_1254F | Forward | GCTGTGGGTAAAGAATTCAAC |
| | HA4_1712R | Reverse | CTGCAAAGACCCATTAGAAC |

**Table S2. Number of families with two or more members used in single-site error frequency calculation**

| Amplicon | Number of families | | | |
|---|---|---|---|---|
| | mRNA-(1) | mRNA-(2) | vRNA-(1) | vRNA-(2) |
| HA1 | 817816 | 459558 | 273944 | 371179 |
| HA2 | 842374 | 469693 | 251809 | 313410 |
| HA3 | 788021 | 615020 | 65088 | 27537 |
| HA4 | 935357 | 450205 | 279034 | 509287 |

**Table S3. Results of SafeSeqS sequencing data for the mRNA-(1) sample.** This Table is included in a separate Excel file (TablesS3S4S5S6.xlsl).

**Table S4. Results of SafeSeqS sequencing data for the mRNA-(2) sample.** This Table is included in a separate Excel file (TablesS3S4S5S6.xlsl).

**Table S5. Results of SafeSeqS sequencing data for the vRNA-(1) sample.** This Table is included in a separate Excel file (TablesS3S4S5S6.xlsl).

**Table S6. Results of SafeSeqS sequencing data for the vRNA-(2) sample.** This Table is included in a separate Excel file (TablesS3S4S5S6.xlsl).

**Table S7. Error frequencies determined for nucleotide changes determined twice, as they correspond to positions at overlapping amplicon regions.** This Table is provided as a separate Excel file (TableS7.xlsl).

**Table S8. Average frequencies for single-site nucleotide substitutions in the error landscape of hemagglutinin-encoding mRNA.** The average of the frequencies obtained for the mRNA-(1) and mRNA-(2) samples. The frequency range defined by the two values being averaged is also included. This Table is provided as a separate Excel file (TableS8.xlsl).

**Table S9. Error-frequency metrics for amino acid replacements in the hemagglutinin molecule.** A metric of the propensity of each amino acid replacement is calculated as the sum of the error frequencies (from Table S8) for all its encoding single-nucleotide changes. The table is provided as a separate Excel file (TableS9.xlsl).

**Table S10. Nucleotide changes for the hemagglutinin gene from natural strains with respect to A/WSN/1933.** The nucleotide changes were identified from the amino-acid-based alignments (Figure S6) and the corresponding nucleotide sequences. The table classifies the 18 natural strains considered into the three sets. These three sets are defined in the main text, but we note here that set 2 corresponds to the H1 strains recommended by the World Health Organization for inclusion in vaccines for the influenza seasons spanning the period 2019-2025 (https://www.who.int/teams/global-influenza-programme/vaccines). The table is provided as a separate Excel file (TableS10.xlsx).

**Table S11. Laboratory experiments on antibody escape with natural virus strains.** The experiments are described in terms of the literature reference, the virus strain and the antibodies used, as well as the number and specification of the escape mutations. This table is provided as a separate Excel file (TableS11.xlsx).

**Table S12. Antibody-escape mutations identified in deep mutational scanning (DMS) experiments on the A/WSN/1933 strain.** Antibody-escape mutations (*i.e.,* amino acid replacements) reported in DMS experiments on the A/WSN/1933 strain (Doud et al., 2017; Doud et al. 2018) are listed, together with virus survival fraction enabled by the mutation, the corresponding metric for the propensity of the mutation according to the error landscape (Table S9) and the antibody used in the DMS experiment. Two sets of data are provided: mutations with survival fraction higher than 0.2 and mutations with survival fraction higher than 0.05. This table is provided as a separate Excel file (TableS12.xlsl).

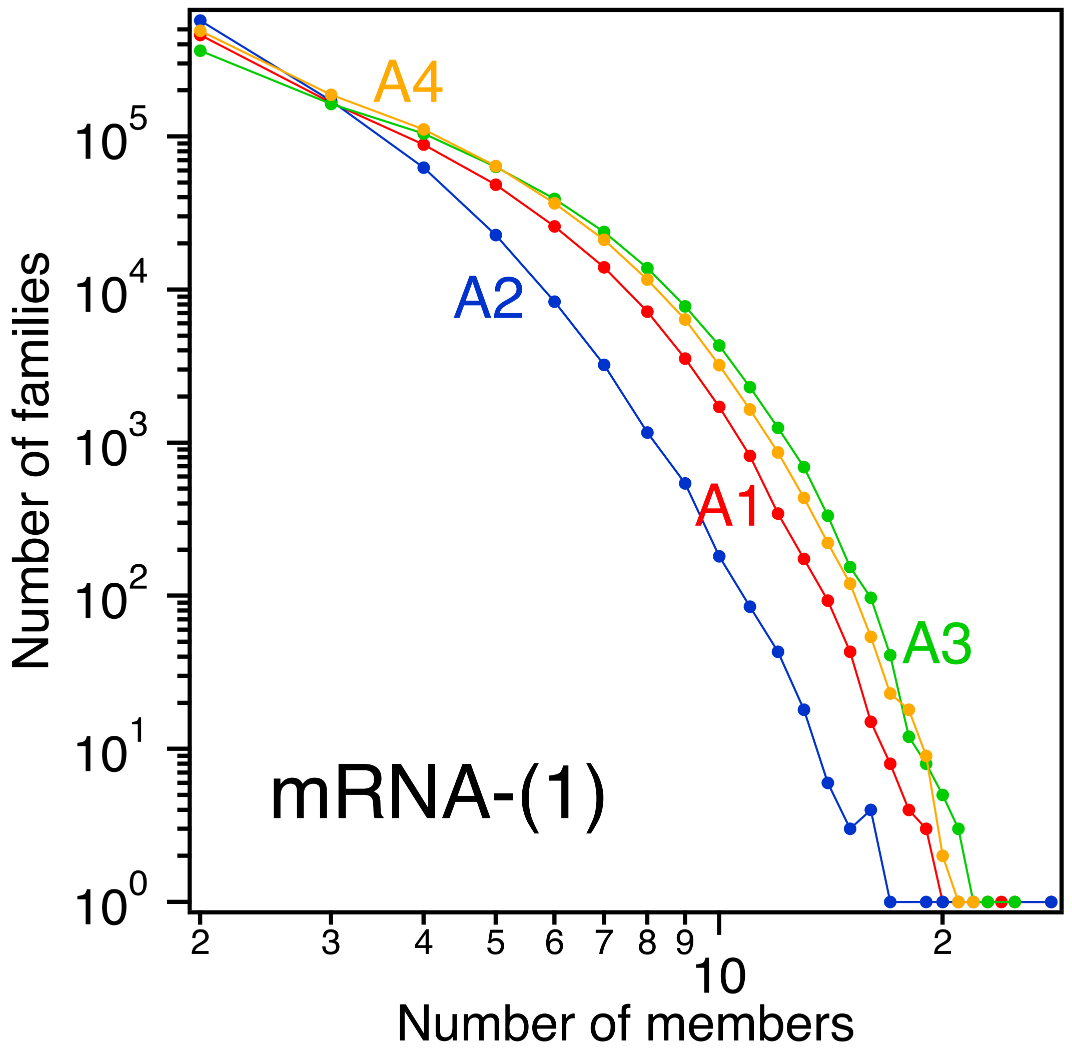


**Figure S1. UMI family distributions for the Safe-Seqs sequencing of the mRNA-(1) biological sample.** Profiles of number of families versus number of members per family are shown for the four amplicons. Note the logarithmic scale for both axes.

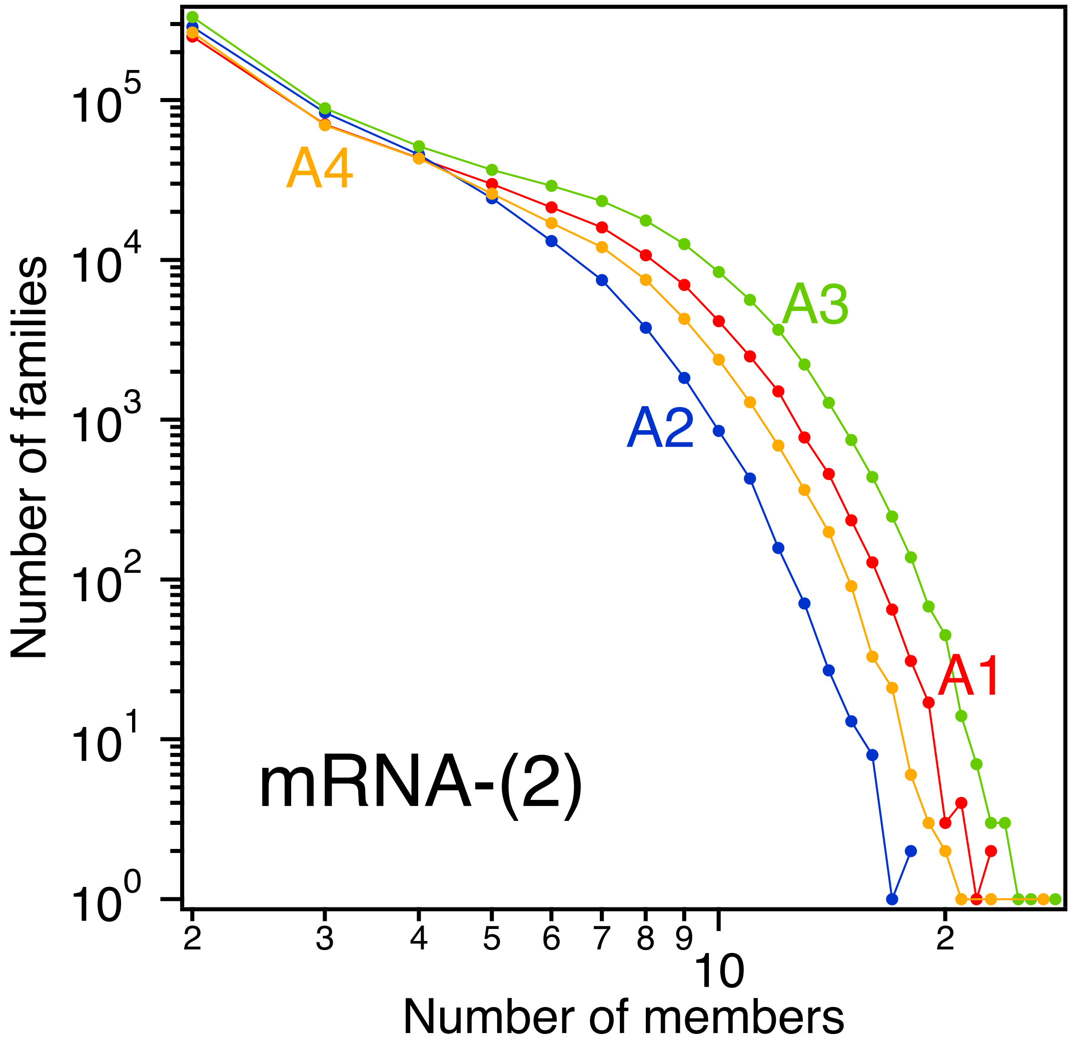


**Figure S2. UMI family distributions for the Safe-Seqs sequencing of the mRNA-(2) biological sample.** Profiles of number of families versus number of members per family are shown for the four amplicons. Note the logarithmic scale for both axes.

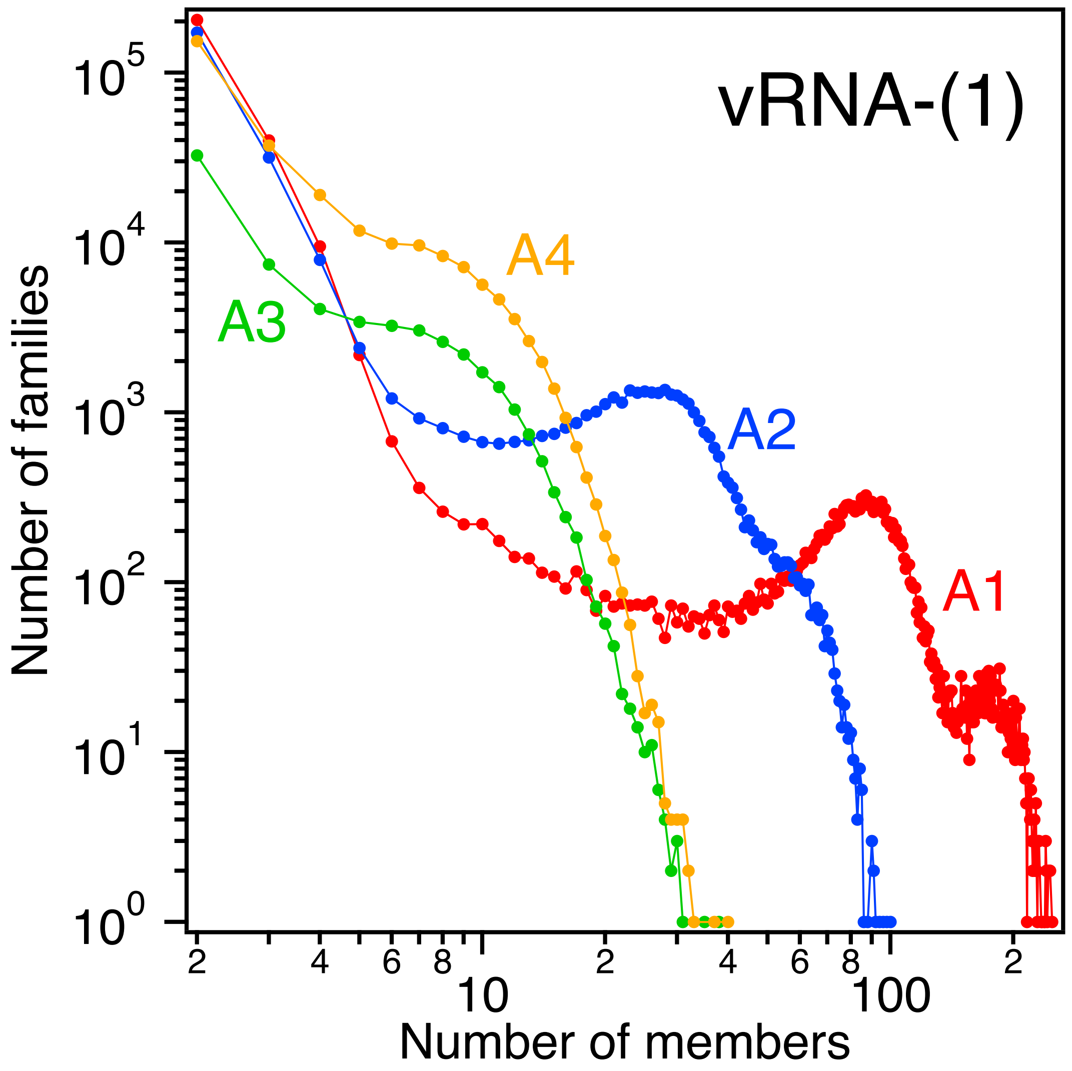


**Figure S3. UMI family distributions for the Safe-Seqs sequencing of the vRNA-(1) biological sample.** Profiles of number of families versus number of members per family are shown for the four amplicons. Note the logarithmic scale for both axes.

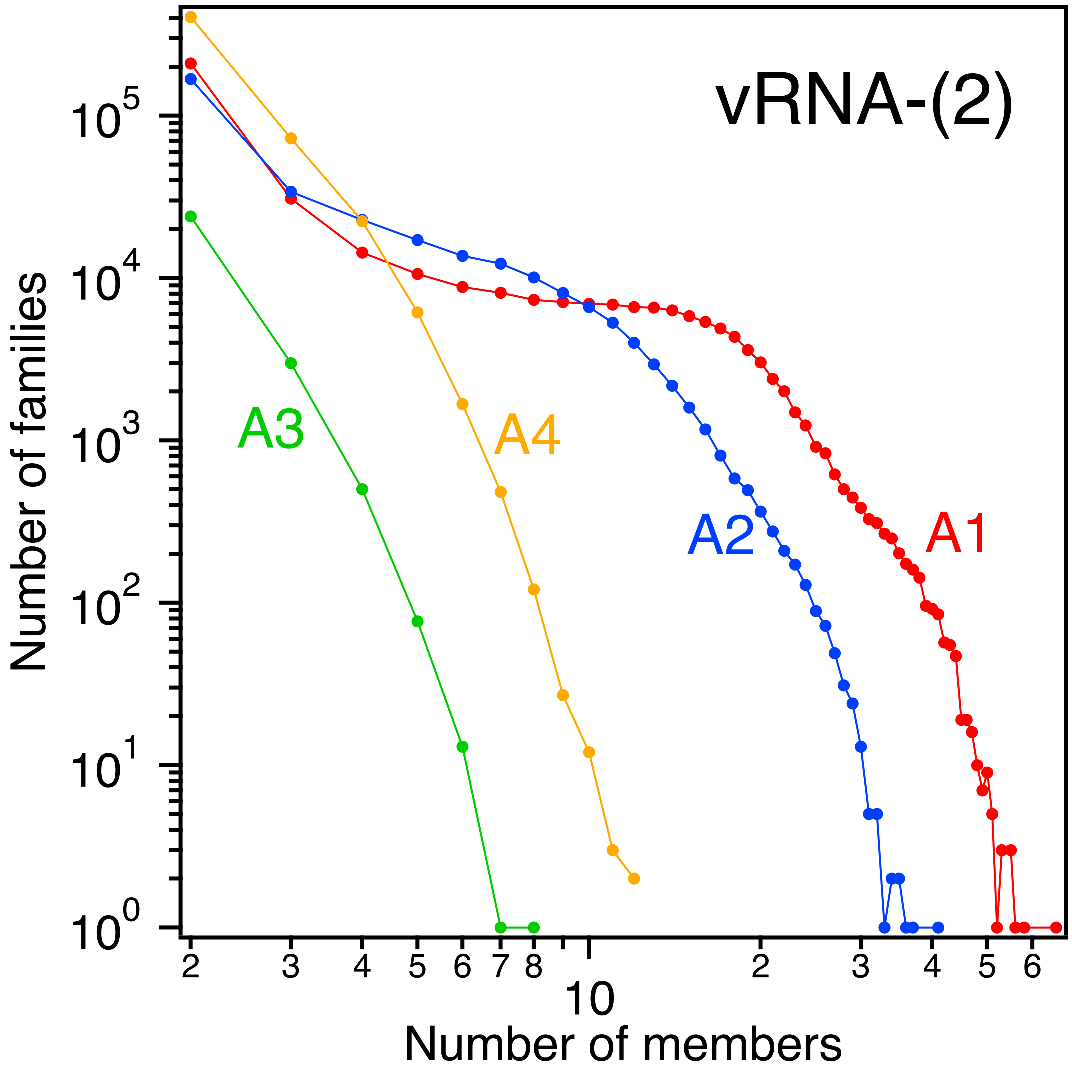


**Figure S4. UMI family distributions for the Safe-Seqs sequencing of the vRNA-(2) biological sample.** Profiles of number of families versus number of members per family are shown for the four amplicons. Note the logarithmic scale for both axes.

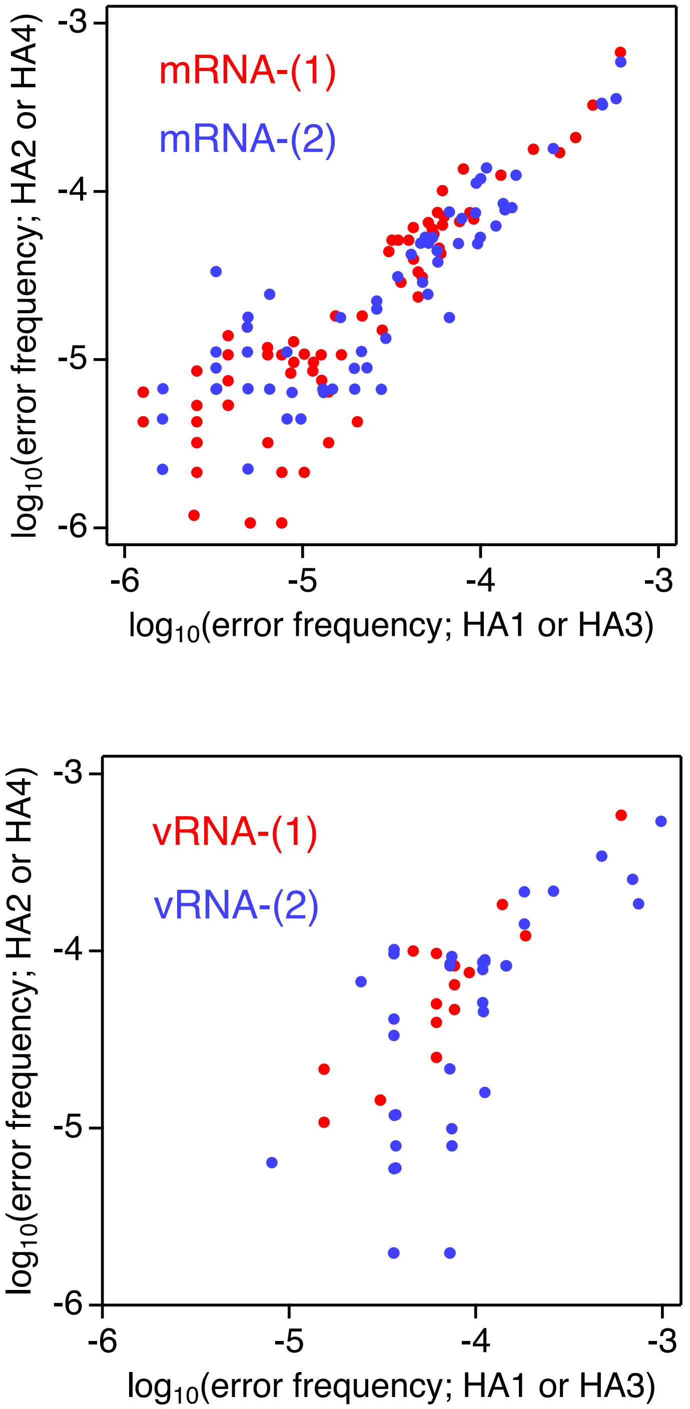


**Figure S5. Correlation between error frequencies determined from different amplicons.** Amplicons HA2 and HA4, as well as amplicons HA3 and HA4, overlap to some extent. For a few errors at positions in overlapping amplicon regions, frequencies are determined twice for each of the biological mRNA or vRNA samples (Table S7). The plots show the frequency of a given error as determined in the HA2 or HA4 amplicons versus the frequency of the same error as determined in HA1 or HA3 amplicons, respectively.

#### A/WSN/1933 vs A/Denver/1957

```
A/WSN/1933         1 MKAKLLVLLYAFVATDADTICIGYHANNSTDTVDTILEKNVAVTHSVNLL     50
                     ||||||:||.|..||||||||||||||||||||||:|||||.||||||||
A/Denver/1957      1 MKAKLLILLCALSATDADTICIGYHANNSTDTVDTVLEKNVTVTHSVNLL     50

A/WSN/1933        51 EDSHNGKLCKLKGIAPLQLGKCNITGWLLGNPECDSLLPARSWSYIVETP    100
                     |||||||||:|||.||||||.|||.||:||||||:|||..||||||.|||
A/Denver/1957     51 EDSHNGKLCRLKGKAPLQLGNCNIAGWVLGNPECESLLSNRSWSYIAETP    100

A/WSN/1933       101 NSENGACYPGDLIDYEELREQLSSVSSLERFEIFPKESSWPNHTFNGVTV    150
                     |||||.|||||..||||||||||||||.|||||||||.||||||..|||.
A/Denver/1957    101 NSENGTCYPGDFADYEELREQLSSVSSFERFEIFPKERSWPNHTTRGVTA    150

A/WSN/1933       151 SCSHRGKSSFYRNLLWLTKKGDSYPKLTNSYVNNKGKEVLVLWGVHHPSS    200
                     :|.|..|||||:||:|||:...|||.|:.|||||:.|||||||||||||:
A/Denver/1957    151 ACPHARKSSFYKNLVWLTEANGSYPNLSRSYVNNQEKEVLVLWGVHHPSN    200

A/WSN/1933       201 SDEQQSLYSNGNAYVSVASSNYNRRFTPEIAARPKVRDQHGRMNYYWTLL    250
                     .:||::||...||||||.|||||||||||||.|||||||.||||||||||
A/Denver/1957    201 IEEQRALYRKDNAYVSVVSSNYNRRFTPEIAKRPKVRDQSGRMNYYWTLL    250

A/WSN/1933       251 EPGDTIIFEATGNLIAPWYAFALSRGFESGIITSNASMHECNTKCQTPQG    300
                     ||||||||||||||||||||||||||..||||||||.:.||:||||||||
A/Denver/1957    251 EPGDTIIFEATGNLIAPWYAFALSRGPGSGIITSNAPLDECDTKCQTPQG    300

A/WSN/1933       301 AINSNLPFQNIHPVTIGECPKYVRSTKLRMVTGLRNIPSIQYRGLFGAIA    350
                     ||||:||||||||||||||||||||||||||||||||||:|.||||||||
A/Denver/1957    301 AINSSLPFQNIHPVTIGECPKYVRSTKLRMVTGLRNIPSVQSRGLFGAIA    350

A/WSN/1933       351 GFIEGGWTGMIDGWYGYHHQNEQGSGYAADQKSTQNAINGITNKVNSVIE    400
                     ||||||||||:|||||||||||||||||||||||||||||||||||||||
A/Denver/1957    351 GFIEGGWTGMMDGWYGYHHQNEQGSGYAADQKSTQNAINGITNKVNSVIE    400

A/WSN/1933       401 KMNTQFTAVGKEFNNLEKRMENLNKKVDDGFLDIWTYNAELLVLLENERT    450
                     ||||||||||||||.||||||||||||||||:||||||||||||||||||
A/Denver/1957    401 KMNTQFTAVGKEFNKLEKRMENLNKKVDDGFMDIWTYNAELLVLLENERT    450

A/WSN/1933       451 LDFHDLNVKNLYEKVKSQLKNNAKEIGNGCFEFYHKCDNECMESVRNGTY    500
                     |||||.||||||||||:||:|||||:|||||||||||||||||||:||||
A/Denver/1957    451 LDFHDSNVKNLYEKVKNQLRNNAKELGNGCFEFYHKCDNECMESVKNGTY    500

A/WSN/1933       501 DYPKYSEESKLNREKIDGVKLESMGVYQILAIYSTVASSLVLLVSLGAIS    550
                     |||||||||||||||||||||||||||:||||||||||||||||||||||
A/Denver/1957    501 DYPKYSEESKLNREKIDGVKLESMGVYRILAIYSTVASSLVLLVSLGAIS    550

A/WSN/1933       551 FWMCSNGSLQCRICI    565
                     |||||||||||||||
A/Denver/1957    551 FWMCSNGSLQCRICI    565
```

#### A/WSN/1933 vs A/USSR/90/1977

```
A/WSN/1933         1 MKAKLLVLLYAFVATDADTICIGYHANNSTDTVDTILEKNVAVTHSVNLL     50
                     |||||||||.|..||||||||||||||||||||||:|||||.||||||||
A/USSR/90/197      1 MKAKLLVLLCALSATDADTICIGYHANNSTDTVDTVLEKNVTVTHSVNLL     50

A/WSN/1933        51 EDSHNGKLCKLKGIAPLQLGKCNITGWLLGNPECDSLLPARSWSYIVETP    100
                     |||||||||:||||||||||||||.||:||||||:||...:|||||.|||
A/USSR/90/197     51 EDSHNGKLCRLKGIAPLQLGKCNIAGWILGNPECESLFSKKSWSYIAETP    100

A/WSN/1933       101 NSENGACYPGDLIDYEELREQLSSVSSLERFEIFPKESSWPNHTFNGVTV    150
                     |||||.||||...||||||||||||||.|||||||||.|||.|...|||.
A/USSR/90/197    101 NSENGTCYPGYFADYEELREQLSSVSSFERFEIFPKERSWPKHNVRGVTA    150

A/WSN/1933       151 SCSHRGKSSFYRNLLWLTKKGDSYPKLTNSYVNNKGKEVLVLWGVHHPSS    200
                     ||||:|||||||||||||:|..|||.|:.||||||.|||||||||||||:
A/USSR/90/197    151 SCSHKGKSSFYRNLLWLTEKNGSYPNLSKSYVNNKEKEVLVLWGVHHPSN    200

A/WSN/1933       201 SDEQQSLYSNGNAYVSVASSNYNRRFTPEIAARPKVRDQHGRMNYYWTLL    250
                     .::|:::|...||||||.|||||||||||||.|||||.|.||:|||||||
A/USSR/90/197    201 IEDQKTIYRKENAYVSVVSSNYNRRFTPEIAERPKVRGQAGRINYYWTLL    250
```

```
A/WSN/1933        251 EPGDTIIFEATGNLIAPWYAFALSRGFESGIITSNASMHECNTKCQTPQG    300
                      ||||||||||.|||||||:||||:|||.||||||||||.||:||||||||
A/USSR/90/197     251 EPGDTIIFEANGNLIAPWHAFALNRGFGSGIITSNASMDECDTKCQTPQG    300

A/WSN/1933        301 AINSNLPFQNIHPVTIGECPKYVRSTKLRMVTGLRNIPSIQYRGLFGAIA    350
                      ||||:||||||||||||||||||||||||||||||||||||.||||||||
A/USSR/90/197     301 AINSSLPFQNIHPVTIGECPKYVRSTKLRMVTGLRNIPSIQSRGLFGAIA    350

A/WSN/1933        351 GFIEGGWTGMIDGWYGYHHQNEQGSGYAADQKSTQNAINGITNKVNSVIE    400
                      ||||||||||||||||||||||||||||||||||||||||||||||||||
A/USSR/90/197     351 GFIEGGWTGMIDGWYGYHHQNEQGSGYAADQKSTQNAINGITNKVNSVIE    400

A/WSN/1933        401 KMNTQFTAVGKEFNNLEKRMENLNKKVDDGFLDIWTYNAELLVLLENERT    450
                      ||||||||||||||.|||||||||||||||||||||||||||||||||||
A/USSR/90/197     401 KMNTQFTAVGKEFNKLEKRMENLNKKVDDGFLDIWTYNAELLVLLENERT    450

A/WSN/1933        451 LDFHDLNVKNLYEKVKSQLKNNAKEIGNGCFEFYHKCDNECMESVRNGTY    500
                      |||||.|||||||||||||||||||||||||||:||||||||:|||||||
A/USSR/90/197     451 LDFHDSNVKNLYEKVKSQLKNNAKEIGNGCFEFYHKCNNECMESVKNGTY    500

A/WSN/1933        501 DYPKYSEESKLNREKIDGVKLESMGVYQILAIYSTVASSLVLLVSLGAIS    550
                      ||||||||||||||||||||||||||||||||||||||||||||||||||
A/USSR/90/197     501 DYPKYSEESKLNREKIDGVKLESMGVYQILAIYSTVASSLVLLVSLGAIS    550

A/WSN/1933        551 FWMCSNGSLQCRICI    565
                      |||||||||||||||
A/USSR/90/197     551 FWMCSNGSLQCRICI    565
```

#### A/WSN/1933 vs A/Memphis/2/1996

```
A/WSN/1933          1 MKAKLLVLLYAFVATDADTICIGYHANNSTDTVDTILEKNVAVTHSVNLL     50
                      ||.||||||.||.||.||||||||||||||||||||:|||||.|||||||
A/Memphis/2/1       1 MKVKLLVLLCAFTATYADTICIGYHANNSTDTVDTVLEKNVTVTHSVNLL     50

A/WSN/1933         51 EDSHNGKLCKLKGIAPLQLGKCNITGWLLGNPECDSLLPARSWSYIVETP    100
                      |||||||||:|||.||||||.|::.||:||||||:||....|||||.|||
A/Memphis/2/1      51 EDSHNGKLCRLKGTAPLQLGNCSVAGWILGNPECESLFSKESWSYIAETP    100

A/WSN/1933        101 NSENGACYPGDLIDYEELREQLSSVSSLERFEIFPKESSWPNHTFNGVTV    150
                      |.|||.||||...|||||||||||||||.|||||||||||||||..|||.
A/Memphis/2/1     101 NPENGTCYPGYFADYEELREQLSSVSSFERFEIFPKESSWPNHTVKGVTA    150

A/WSN/1933        151 SCSHRGKSSFYRNLLWLTKKGDSYPKLTNSYVNNKGKEVLVLWGVHHPSS    200
                      ||||.||||||:||||||:|...||.|:.||||||.|||||||||||||:
A/Memphis/2/1     151 SCSHNGKSSFYKNLLWLTEKNGLYPNLSKSYVNNKEKEVLVLWGVHHPSN    200

A/WSN/1933        201 SDEQQSLYSNGNAYVSVASSNYNRRFTPEIAARPKVRDQHGRMNYYWTLL    250
                      ..:|:::|...||||||.||:|:||||||||.|||||||.||:|||||||
A/Memphis/2/1     201 IGDQRAIYHTENAYVSVVSSHYSRRFTPEIAKRPKVRDQEGRINYYWTLL    250

A/WSN/1933        251 EPGDTIIFEATGNLIAPWYAFALSRGFESGIITSNASMHECNTKCQTPQG    300
                      ||||||||||.|||||||||||||||||.|||||||||.||:.|||||||
A/Memphis/2/1     251 EPGDTIIFEANGNLIAPWYAFALSRGFGSGIITSNASMGECDAKCQTPQG    300

A/WSN/1933        301 AINSNLPFQNIHPVTIGECPKYVRSTKLRMVTGLRNIPSIQYRGLFGAIA    350
                      ||||:|||||:||||||||||||||||||||||||||||||.||||||||
A/Memphis/2/1     301 AINSSLPFQNVHPVTIGECPKYVRSTKLRMVTGLRNIPSIQSRGLFGAIA    350

A/WSN/1933        351 GFIEGGWTGMIDGWYGYHHQNEQGSGYAADQKSTQNAINGITNKVNSVIE    400
                      ||||||||||||||||||||||||||||||||||||||:|||||||||||
A/Memphis/2/1     351 GFIEGGWTGMIDGWYGYHHQNEQGSGYAADQKSTQNAIDGITNKVNSVIE    400

A/WSN/1933        401 KMNTQFTAVGKEFNNLEKRMENLNKKVDDGFLDIWTYNAELLVLLENERT    450
                      ||||||||||||||.||:||||||||||||||||||||||||||||||||
A/Memphis/2/1     401 KMNTQFTAVGKEFNKLERRMENLNKKVDDGFLDIWTYNAELLVLLENERT    450

A/WSN/1933        451 LDFHDLNVKNLYEKVKSQLKNNAKEIGNGCFEFYHKCDNECMESVRNGTY    500
                      |||||.||||||||||:||||||||||||||||:||||||||:|||||||
A/Memphis/2/1     451 LDFHDSNVKNLYEKVKNQLKNNAKEIGNGCFEFYHKCNNECMESVKNGTY    500

A/WSN/1933        501 DYPKYSEESKLNREKIDGVKLESMGVYQILAIYSTVASSLVLLVSLGAIS    550
                      ||||||||||||||||||||||||||||||||||||||||||||||||||
A/Memphis/2/1     501 DYPKYSEESKLNREKIDGVKLESMGVYQILAIYSTVASSLVLLVSLGAIS    550
```

```
A/WSN/1933       551 FWMCSNGSLQCRICI    565
                     |||||||||||||||
A/Memphis/2/1    551 FWMCSNGSLQCRICI    565
```

### A/WSN/1933 vs A/Managua/156.01/2008

```
A/WSN/1933         1 MKAKLLVLLYAFVATDADTICIGYHANNSTDTVDTILEKNVAVTHSVNLL     50
                     ||.||||||..|.||.|||||||||||||||||||:|||||.||||||||
A/Managua/156      1 MKVKLLVLLCTFTATYADTICIGYHANNSTDTVDTVLEKNVTVTHSVNLL     50

A/WSN/1933        51 EDSHNGKLCKLKGIAPLQLGKCNITGWLLGNPECDSLLPARSWSYIVETP    100
                     |:|||||||.||||||||||.|::.||:||||||:.|:...|||||||.|
A/Managua/156     51 ENSHNGKLCLLKGIAPLQLGNCSVAGWILGNPECELLISKESWSYIVEKP    100

A/WSN/1933       101 NSENGACYPGDLIDYEELREQLSSVSSLERFEIFPKESSWPNHTFNGVTV    150
                     |.|||.||||...||||||||||||||.|||||||||.||||||..||:.
A/Managua/156    101 NPENGTCYPGHFADYEELREQLSSVSSFERFEIFPKEGSWPNHTVTGVSA    150

A/WSN/1933       151 SCSHRGKSSFYRNLLWLTKKGDSYPKLTNSYVNNKGKEVLVLWGVHHPSS    200
                     ||||.|:|||||||||||.|...||.|:.||.|||.||||||||||||.:
A/Managua/156    151 SCSHNGESSFYRNLLWLTGKNGLYPNLSKSYANNKEKEVLVLWGVHHPPN    200

A/WSN/1933       201 SDEQQSLYSNGNAYVSVASSNYNRRFTPEIAARPKVRDQHGRMNYYWTLL    250
                     ..:|::||...||||||.||:|:|:||||||.|||||||.||:|||||||
A/Managua/156    201 IGDQKALYHTENAYVSVVSSHYSRKFTPEIAKRPKVRDQEGRINYYWTLL    250

A/WSN/1933       251 EPGDTIIFEATGNLIAPWYAFALSRGFESGIITSNASMHECNTKCQTPQG    300
                     ||||||||||.||||||.|||||||||.||||.|||.|.:|:.|||||||
A/Managua/156    251 EPGDTIIFEANGNLIAPRYAFALSRGFGSGIINSNAPMDKCDAKCQTPQG    300

A/WSN/1933       301 AINSNLPFQNIHPVTIGECPKYVRSTKLRMVTGLRNIPSIQYRGLFGAIA    350
                     ||||:|||||:||||||||||||||.|||||||||||||||.||||||||
A/Managua/156    301 AINSSLPFQNVHPVTIGECPKYVRSAKLRMVTGLRNIPSIQSRGLFGAIA    350

A/WSN/1933       351 GFIEGGWTGMIDGWYGYHHQNEQGSGYAADQKSTQNAINGITNKVNSVIE    400
                     ||||||||||:|||||||||||||||||||||||||||||||||||||||
A/Managua/156    351 GFIEGGWTGMVDGWYGYHHQNEQGSGYAADQKSTQNAINGITNKVNSVIE    400

A/WSN/1933       401 KMNTQFTAVGKEFNNLEKRMENLNKKVDDGFLDIWTYNAELLVLLENERT    450
                     ||||||||||||||.||:|||||||||||||:||||||||||||||||||
A/Managua/156    401 KMNTQFTAVGKEFNKLERRMENLNKKVDDGFIDIWTYNAELLVLLENERT    450

A/WSN/1933       451 LDFHDLNVKNLYEKVKSQLKNNAKEIGNGCFEFYHKCDNECMESVRNGTY    500
                     |||||.|||||||||||||||||||||||||||||||::||||||:||||
A/Managua/156    451 LDFHDSNVKNLYEKVKSQLKNNAKEIGNGCFEFYHKCNDECMESVKNGTY    500

A/WSN/1933       501 DYPKYSEESKLNREKIDGVKLESMGVYQILAIYSTVASSLVLLVSLGAIS    550
                     ||||||||||||||||||||||||||||||||||||||||||||||||||
A/Managua/156    501 DYPKYSEESKLNREKIDGVKLESMGVYQILAIYSTVASSLVLLVSLGAIS    550

A/WSN/1933       551 FWMCSNGSLQCRICI    565
                     |||||||||||||||
A/Managua/156    551 FWMCSNGSLQCRICI    565
```

### A/WSN/1933 vs A/Texas/24/2012

```
A_WSN_1933         1 MKAKLLVLLYAFVATDADTICIGYHANNSTDTVDTILEKNVAVTHSVNLL     50
                     |||.|:||||.|...:|||:|||||||||||||||:|||||.||||||||
A_Texas_24_20      1 MKAILVVLLYTFATANADTLCIGYHANNSTDTVDTVLEKNVTVTHSVNLL     50

A_WSN_1933        51 EDSHNGKLCKLKGIAPLQLGKCNITGWLLGNPECDSLLPARSWSYIVETP    100
                     ||.||||||||:|:|||.||||||.||:||||||::|..|.||||||||.
A_Texas_24_20     51 EDKHNGKLCKLRGVAPLHLGKCNIAGWILGNPECETLSTASSWSYIVETS    100

A_WSN_1933       101 NSENGACYPGDLIDYEELREQLSSVSSLERFEIFPKESSWPNHTFNGVTV    150
                     :|:||.|||||.|||||||||||||||.||||||||.||||||..||||.
A_Texas_24_20    101 SSDNGTCYPGDFIDYEELREQLSSVSSFERFEIFPKTSSWPNHDSNGVTA    150
```

```
A_WSN_1933         151 SCSHRGKSSFYRNLLWLTKKGDSYPKLTNSYVNNKGKEVLVLWGVHHPSS    200
                       :|.|.|...||:||:||.|||:|||||:.||:|:||||||||||:||||:
A_Texas_24_20      151 ACPHAGAKGFYKNLIWLVKKGNSYPKLSKSYINDKGKEVLVLWGIHHPST    200

A_WSN_1933         201 SDEQQSLYSNGNAYVSVASSNYNRRFTPEIAARPKVRDQHGRMNYYWTLL    250
                       :.:|||||.|.:.||.|.:|.|:::|.||||.|||||||.|||||||||:
A_Texas_24_20      201 TADQQSLYQNADTYVFVGTSRYSKKFKPEIAIRPKVRDQEGRMNYYWTLV    250

A_WSN_1933         251 EPGDTIIFEATGNLIAPWYAFALSRGFESGIITSNASMHECNTKCQTPQG    300
                       ||||.|.|||||||:.|.||||:.|...||||.|:..:|:|||.||||:|
A_Texas_24_20      251 EPGDKITFEATGNLVVPRYAFAMERDAGSGIIISDTPVHDCNTTCQTPKG    300

A_WSN_1933         301 AINSNLPFQNIHPVTIGECPKYVRSTKLRMVTGLRNIPSIQYRGLFGAIA    350
                       |||::||||||||:|||:|||||:|||||:.|||||:||||.||||||||
A_Texas_24_20      301 AINTSLPFQNIHPITIGKCPKYVKSTKLRLATGLRNVPSIQSRGLFGAIA    350

A_WSN_1933         351 GFIEGGWTGMIDGWYGYHHQNEQGSGYAADQKSTQNAINGITNKVNSVIE    400
                       ||||||||||:|||||||||||||||||||.|||||||:.||||||||||
A_Texas_24_20      351 GFIEGGWTGMVDGWYGYHHQNEQGSGYAADLKSTQNAIDKITNKVNSVIE    400

A_WSN_1933         401 KMNTQFTAVGKEFNNLEKRMENLNKKVDDGFLDIWTYNAELLVLLENERT    450
                       ||||||||||||||:||||:||||||||||||||||||||||||||||||
A_Texas_24_20      401 KMNTQFTAVGKEFNHLEKRIENLNKKVDDGFLDIWTYNAELLVLLENERT    450

A_WSN_1933         451 LDFHDLNVKNLYEKVKSQLKNNAKEIGNGCFEFYHKCDNECMESVRNGTY    500
                       ||:||.|||||||||::||||||||||||||||||||||.|||||:||||
A_Texas_24_20      451 LDYHDSNVKNLYEKVRNQLKNNAKEIGNGCFEFYHKCDNTCMESVKNGTY    500

A_WSN_1933         501 DYPKYSEESKLNREKIDGVKLESMGVYQILAIYSTVASSLVLLVSLGAIS    550
                       ||||||||:|||||:||||||||..:|||||||||.||||||:|||||||
A_Texas_24_20      501 DYPKYSEEAKLNREEIDGVKLESTRIYQILAIYSTAASSLVLVVSLGAIS    550

A_WSN_1933         551 FWMCSNGSLQCRICI    565
                       |||||||||||||||
A_Texas_24_20      551 FWMCSNGSLQCRICI    565
```

#### A/WSN/1933 vs A/Wisconsin/588/2019

```
A/WSN/1933           1 MKAKLLVLLYAFVATDADTICIGYHANNSTDTVDTILEKNVAVTHSVNLL     50
                       |||.|:|:||.|...:|||:|||||||||||||||:|||||.||||||||
A/Wisconsin/5        1 MKAILVVMLYTFTTANADTLCIGYHANNSTDTVDTVLEKNVTVTHSVNLL     50

A/WSN/1933          51 EDSHNGKLCKLKGIAPLQLGKCNITGWLLGNPECDSLLPARSWSYIVETP    100
                       ||.||||||||:|:|||.||||||.||:||||||:||..||||||||||.
A/Wisconsin/5       51 EDKHNGKLCKLRGVAPLHLGKCNIAGWILGNPECESLSTARSWSYIVETS    100

A/WSN/1933         101 NSENGACYPGDLIDYEELREQLSSVSSLERFEIFPKESSWPNH-TFNGVT    149
                       ||:||.|||||.|:|||||||||||||.||||||||.|||||| :.||||
A/Wisconsin/5      101 NSDNGTCYPGDFINYEELREQLSSVSSFERFEIFPKTSSWPNHDSDNGVT    150

A/WSN/1933         150 VSCSHRGKSSFYRNLLWLTKKGDSYPKLTNSYVNNKGKEVLVLWGVHHPS    199
                       .:|.|.|..|||:||:||.|||.||||:..:|:|:||||||||||:|||.
A/Wisconsin/5      151 AACPHAGAKSFYKNLIWLVKKGKSYPKINQTYINDKGKEVLVLWGIHHPP    200

A/WSN/1933         200 SSDEQQSLYSNGNAYVSVASSNYNRRFTPEIAARPKVRDQHGRMNYYWTL    249
                       :..:|||||.|.:|||.|.:|.|:::|.||||.|||||||.|||||||||
A/Wisconsin/5      201 TIADQQSLYQNADAYVFVGTSRYSKKFKPEIATRPKVRDQEGRMNYYWTL    250

A/WSN/1933         250 LEPGDTIIFEATGNLIAPWYAFALSRGFESGIITSNASMHECNTKCQTPQ    299
                       :||||.|.|||||||:||.|||.:.|...||||.|:..:|:|||.||||:
A/Wisconsin/5      251 VEPGDKITFEATGNLVAPRYAFTMERDAGSGIIISDTPVHDCNTTCQTPE    300

A/WSN/1933         300 GAINSNLPFQNIHPVTIGECPKYVRSTKLRMVTGLRNIPSIQYRGLFGAI    349
                       ||||::|||||:||:|||:|||||:|||||:.|||||:||||.|||||||
A/Wisconsin/5      301 GAINTSLPFQNVHPITIGKCPKYVKSTKLRLATGLRNVPSIQSRGLFGAI    350

A/WSN/1933         350 AGFIEGGWTGMIDGWYGYHHQNEQGSGYAADQKSTQNAINGITNKVNSVI    399
                       |||||||||||:|||||||||||||||||||.|||||||:.|||||||||
A/Wisconsin/5      351 AGFIEGGWTGMVDGWYGYHHQNEQGSGYAADLKSTQNAIDKITNKVNSVI    400

A/WSN/1933         400 EKMNTQFTAVGKEFNNLEKRMENLNKKVDDGFLDIWTYNAELLVLLENER    449
                       |||||||||||||||:||||:|||||||||||||||||||||||||||||
A/Wisconsin/5      401 EKMNTQFTAVGKEFNHLEKRIENLNKKVDDGFLDIWTYNAELLVLLENER    450
```

```
A/WSN/1933       450 TLDFHDLNVKNLYEKVKSQLKNNAKEIGNGCFEFYHKCDNECMESVRNGT    499
                     |||:||.|||||||||::|||||||||||||||||||||||.|||||:|||
A/Wisconsin/5    451 TLDYHDSNVKNLYEKVRNQLKNNAKEIGNGCFEFYHKCDNTCMESVKNGT    500

A/WSN/1933       500 YDYPKYSEESKLNREKIDGVKLESMGVYQILAIYSTVASSLVLLVSLGAI    549
                     |||||||||:||||||||||||:|..:||||||||||||||||:||||||
A/Wisconsin/5    501 YDYPKYSEEAKLNREKIDGVKLDSTRIYQILAIYSTVASSLVLVVSLGAI    550

A/WSN/1933       550 SFWMCSNGSLQCRICI    565
                     ||||||||||||||||
A/Wisconsin/5    551 SFWMCSNGSLQCRICI    566
```

#### A/WSN/1933 vs A/Victoria/2570/2019

```
A/WSN/1933         1 MKAKLLVLLYAFVATDADTICIGYHANNSTDTVDTILEKNVAVTHSVNLL     50
                     |||.|:|:||.|...:|||:||||||||||||||:|||||.||||||||
A/Victoria/25      1 MKAILVVMLYTFTTANADTLCIGYHANNSTDTVDTVLEKNVTVTHSVNLL     50

A/WSN/1933        51 EDSHNGKLCKLKGIAPLQLGKCNITGWLLGNPECDSLLPARSWSYIVETP    100
                     ||.||||||||:|:|||.||||||.||:||||||:||..||||||||||.
A/Victoria/25     51 EDKHNGKLCKLRGVAPLHLGKCNIAGWILGNPECESLSTARSWSYIVETS    100

A/WSN/1933       101 NSENGACYPGDLIDYEELREQLSSVSSLERFEIFPKESSWPNH-TFNGVT    149
                     ||:||.|||||.|:|||||||||||||.||||||||.|||||| :.||||
A/Victoria/25    101 NSDNGTCYPGDFINYEELREQLSSVSSFERFEIFPKTSSWPNHDSDNGVT    150

A/WSN/1933       150 VSCSHRGKSSFYRNLLWLTKKGDSYPKLTNSYVNNKGKEVLVLWGVHHPS    199
                     .:|.|.|..|||:||:||.|||.||||:..:|:|:||||||||||:|||.
A/Victoria/25    151 AACPHAGAKSFYKNLIWLVKKGKSYPKINQTYINDKGKEVLVLWGIHHPP    200

A/WSN/1933       200 SSDEQQSLYSNGNAYVSVASSNYNRRFTPEIAARPKVRDQHGRMNYYWTL    249
                     :..:|||||.|.:|||.|.:|.|:::|.||||.||||||:.|||||||||
A/Victoria/25    201 TIADQQSLYQNADAYVFVGTSRYSKKFKPEIATRPKVRDREGRMNYYWTL    250

A/WSN/1933       250 LEPGDTIIFEATGNLIAPWYAFALSRGFESGIITSNASMHECNTKCQTPQ    299
                     :||||.|.|||||||:||.|||.:.|...||||.|:..:|:|||.||||:
A/Victoria/25    251 VEPGDKITFEATGNLVAPRYAFTMERDAGSGIIISDTPVHDCNTTCQTPE    300

A/WSN/1933       300 GAINSNLPFQNIHPVTIGECPKYVRSTKLRMVTGLRNIPSIQYRGLFGAI    349
                     ||||::|||||:||:|||:||||||:|||||:.|||||:||||.|||||||
A/Victoria/25    301 GAINTSLPFQNVHPITIGKCPKYVKSTKLRLATGLRNVPSIQSRGLFGAI    350

A/WSN/1933       350 AGFIEGGWTGMIDGWYGYHHQNEQGSGYAADQKSTQNAINGITNKVNSVI    399
                     |||||||||||:|||||||||||||||||||.|||||||:.||||||||||
A/Victoria/25    351 AGFIEGGWTGMVDGWYGYHHQNEQGSGYAADLKSTQNAIDKITNKVNSVI    400

A/WSN/1933       400 EKMNTQFTAVGKEFNNLEKRMENLNKKVDDGFLDIWTYNAELLVLLENER    449
                     ||||||||||||||:||||:||||||||||||||||||||||||||||||
A/Victoria/25    401 EKMNTQFTAVGKEFNHLEKRIENLNKKVDDGFLDIWTYNAELLVLLENER    450

A/WSN/1933       450 TLDFHDLNVKNLYEKVKSQLKNNAKEIGNGCFEFYHKCDNECMESVRNGT    499
                     |||:||.|||||||||::|||||||||||||||||||||||.|||||:|||
A/Victoria/25    451 TLDYHDSNVKNLYEKVRNQLKNNAKEIGNGCFEFYHKCDNTCMESVKNGT    500

A/WSN/1933       500 YDYPKYSEESKLNREKIDGVKLESMGVYQILAIYSTVASSLVLLVSLGAI    549
                     |||||||||:||||||||||||:|..:||||||||||||||||:||||||
A/Victoria/25    501 YDYPKYSEEAKLNREKIDGVKLDSTRIYQILAIYSTVASSLVLVVSLGAI    550

A/WSN/1933       550 SFWMCSNGSLQCRICI    565
                     ||||||||||||||||
A/Victoria/25    551 SFWMCSNGSLQCRICI    566
```

#### A/WSN/1933 vs A/Hawaii/70/2019

```
A/WSN/1933         1 MKAKLLVLLYAFVATDADTICIGYHANNSTDTVDTILEKNVAVTHSVNLL     50
                     |||.|:||||.|...:|||:||||||||||||||:|||||.||||||||
A/Hawaii/70/2      1 MKAILVVLLYTFTTANADTLCIGYHANNSTDTVDTVLEKNVTVTHSVNLL     50

A/WSN/1933        51 EDSHNGKLCKLKGIAPLQLGKCNITGWLLGNPECDSLLPARSWSYIVETP    100
                     ||.||||||||:|:|||.||||||.||:||||||:||..||||||||||.
A/Hawaii/70/2     51 EDKHNGKLCKLRGVAPLHLGKCNIAGWILGNPECESLSTARSWSYIVETS    100
```

```
A/WSN/1933        101 NSENGACYPGDLIDYEELREQLSSVSSLERFEIFPKESSWPNH-TFNGVT    149
                      ||:||.|||||.|:|||||||||||||.||||||||.|||||| :..|||
A/Hawaii/70/2     101 NSDNGTCYPGDFINYEELREQLSSVSSFERFEIFPKTSSWPNHDSDKGVT    150

A/WSN/1933        150 VSCSHRGKSSFYRNLLWLTKKGDSYPKLTNSYVNNKGKEVLVLWGVHHPS    199
                      .:|.|.|..|||:||:||.|||:|||||..:|:|:||||||||||:|||.
A/Hawaii/70/2     151 AACPHAGAKSFYKNLIWLVKKGNSYPKLNQTYINDKGKEVLVLWGIHHPP    200

A/WSN/1933        200 SSDEQQSLYSNGNAYVSVASSNYNRRFTPEIAARPKVRDQHGRMNYYWTL    249
                      :...|:|||.|.:|||.|.:|.|:::|.||||.|||||||.|||||||||
A/Hawaii/70/2     201 TIAAQESLYQNADAYVFVGTSRYSKKFKPEIATRPKVRDQEGRMNYYWTL    250

A/WSN/1933        250 LEPGDTIIFEATGNLIAPWYAFALSRGFESGIITSNASMHECNTKCQTPQ    299
                      :||||.|.|||||||:.|.|||.:.|...||||.|:..:|:|||.||||:
A/Hawaii/70/2     251 VEPGDKITFEATGNLVVPRYAFTMERDAGSGIIISDTPVHDCNTTCQTPE    300

A/WSN/1933        300 GAINSNLPFQNIHPVTIGECPKYVRSTKLRMVTGLRNIPSIQYRGLFGAI    349
                      ||||::|||||:||:|||:|||||:|||||:.|||||:||||.|||||||
A/Hawaii/70/2     301 GAINTSLPFQNVHPITIGKCPKYVKSTKLRLATGLRNVPSIQSRGLFGAI    350

A/WSN/1933        350 AGFIEGGWTGMIDGWYGYHHQNEQGSGYAADQKSTQNAINGITNKVNSVI    399
                      |||||||||||:|||||||||||||||||||.|||||||:.|||||||||
A/Hawaii/70/2     351 AGFIEGGWTGMVDGWYGYHHQNEQGSGYAADLKSTQNAIDKITNKVNSVI    400

A/WSN/1933        400 EKMNTQFTAVGKEFNNLEKRMENLNKKVDDGFLDIWTYNAELLVLLENER    449
                      |||||||||||||||:||||:|||||||||||||||||||||||||||||
A/Hawaii/70/2     401 EKMNTQFTAVGKEFNHLEKRIENLNKKVDDGFLDIWTYNAELLVLLENER    450

A/WSN/1933        450 TLDFHDLNVKNLYEKVKSQLKNNAKEIGNGCFEFYHKCDNECMESVRNGT    499
                      |||:||.|||||||||::||||||||||||||||||||||.|||||:|||
A/Hawaii/70/2     451 TLDYHDSNVKNLYEKVRNQLKNNAKEIGNGCFEFYHKCDNTCMESVKNGT    500

A/WSN/1933        500 YDYPKYSEESKLNREKIDGVKLESMGVYQILAIYSTVASSLVLLVSLGAI    549
                      |||||||||:||||||||||||||..:||||||||||||||||:||||||
A/Hawaii/70/2     501 YDYPKYSEEAKLNREKIDGVKLESTRIYQILAIYSTVASSLVLVVSLGAI    550

A/WSN/1933        550 SFWMCSNGSLQCRICI    565
                      ||||||||||||||||
A/Hawaii/70/2     551 SFWMCSNGSLQCRICI    566
```

#### A/WSN/1933 vs A/Sydney/5/2021

```
A/WSN/1933          1 MKAKLLVLLYAFVATDADTICIGYHANNSTDTVDTILEKNVAVTHSVNLL     50
                      |||.|:|:||.|...:|||:|||||||||||||||:|||||.||||||||
A/Sydney/5/20       1 MKAILVVMLYTFTTANADTLCIGYHANNSTDTVDTVLEKNVTVTHSVNLL     50

A/WSN/1933         51 EDSHNGKLCKLKGIAPLQLGKCNITGWLLGNPECDSLLPARSWSYIVETP    100
                      ||.||||||||:|:|||.||:|||.||:||||||:||..||||||||||.
A/Sydney/5/20      51 EDKHNGKLCKLRGVAPLHLGQCNIAGWILGNPECESLSTARSWSYIVETS    100

A/WSN/1933        101 NSENGACYPGDLIDYEELREQLSSVSSLERFEIFPKESSWPNH-TFNGVT    149
                      ||:||.||||:.|:|||||||||||||.||||||||.|||||| :.||||
A/Sydney/5/20     101 NSDNGTCYPGNFINYEELREQLSSVSSFERFEIFPKTSSWPNHDSDNGVT    150

A/WSN/1933        150 VSCSHRGKSSFYRNLLWLTKKGDSYPKLTNSYVNNKGKEVLVLWGVHHPS    199
                      .:|.|.|..|||:||:||.|||.||||:..:|:|:||||||||||:|||.
A/Sydney/5/20     151 AACPHAGAKSFYKNLIWLVKKGKSYPKINQTYINDKGKEVLVLWGIHHPP    200

A/WSN/1933        200 SSDEQQSLYSNGNAYVSVASSNYNRRFTPEIAARPKVRDQHGRMNYYWTL    249
                      :..:|:|||.|.:|||.|.:|.|:::|.|||||||||||:.|||||||||
A/Sydney/5/20     201 TITDQESLYQNADAYVFVGTSRYSKKFKPEIAARPKVRDRAGRMNYYWTL    250

A/WSN/1933        250 LEPGDTIIFEATGNLIAPWYAFALSRGFESGIITSNASMHECNTKCQTPQ    299
                      :||||.|.|||||||:||.|||.:.:...||||.|:..:|:|||.||||:
A/Sydney/5/20     251 VEPGDKITFEATGNLVAPRYAFTMEKDAGSGIIISDTPVHDCNTTCQTPE    300

A/WSN/1933        300 GAINSNLPFQNIHPVTIGECPKYVRSTKLRMVTGLRNIPSIQYRGLFGAI    349
                      ||||::|||||:||:|||:|||||||||||:.|||||:||||.|||||||
A/Sydney/5/20     301 GAINTSLPFQNVHPITIGKCPKYVRSTKLRLATGLRNVPSIQSRGLFGAI    350
```

```
A/WSN/1933       350 AGFIEGGWTGMIDGWYGYHHQNEQGSGYAADQKSTQNAINGITNKVNSVI    399
                     |||||||||||:|||||||||||||||||||.|||||||:.|||||||||
A/Sydney/5/20    351 AGFIEGGWTGMVDGWYGYHHQNEQGSGYAADLKSTQNAIDKITNKVNSVI    400

A/WSN/1933       400 EKMNTQFTAVGKEFNNLEKRMENLNKKVDDGFLDIWTYNAELLVLLENER    449
                     |||||||||||||||:||||:|||||||||||||||||||||||||||||
A/Sydney/5/20    401 EKMNTQFTAVGKEFNHLEKRIENLNKKVDDGFLDIWTYNAELLVLLENER    450

A/WSN/1933       450 TLDFHDLNVKNLYEKVKSQLKNNAKEIGNGCFEFYHKCDNECMESVRNGT    499
                     |||:||.|||||||||::||||||||||||||||||||||.|||||:|||
A/Sydney/5/20    451 TLDYHDSNVKNLYEKVRNQLKNNAKEIGNGCFEFYHKCDNTCMESVKNGT    500

A/WSN/1933       500 YDYPKYSEESKLNREKIDGVKLESMGVYQILAIYSTVASSLVLLVSLGAI    549
                     |||||||||:||||||||||||:|..:||||||||||||||||:||||||
A/Sydney/5/20    501 YDYPKYSEEAKLNREKIDGVKLDSTRIYQILAIYSTVASSLVLVVSLGAI    550

A/WSN/1933       550 SFWMCSNGSLQCRICI    565
                     ||||||||||||||||
A/Sydney/5/20    551 SFWMCSNGSLQCRICI    566
```

#### A/WSN/1933 vs A/Victoria/4897/2022

```
A/WSN/1933         1 MKAKLLVLLYAFVATDADTICIGYHANNSTDTVDTILEKNVAVTHSVNLL     50
                     |||.|:|:||.|...:|||:|||||||||||||||:|||||.||||||||
A/Victoria/48      1 MKAILVVMLYTFTTANADTLCIGYHANNSTDTVDTVLEKNVTVTHSVNLL     50

A/WSN/1933        51 EDSHNGKLCKLKGIAPLQLGKCNITGWLLGNPECDSLLPARSWSYIVETP    100
                     ||.||||||||:|:|||.||:|||.||:||||||:||..||||||||||.
A/Victoria/48     51 EDKHNGKLCKLRGVAPLHLGQCNIAGWILGNPECESLSTARSWSYIVETS    100

A/WSN/1933       101 NSENGACYPGDLIDYEELREQLSSVSSLERFEIFPKESSWPNH-TFNGVT    149
                     ||:||.|||||.|:|||||||||||||.||||||||.|||||| :.||||
A/Victoria/48    101 NSDNGTCYPGDFINYEELREQLSSVSSFERFEIFPKTSSWPNHDSDNGVT    150

A/WSN/1933       150 VSCSHRGKSSFYRNLLWLTKKGDSYPKLTNSYVNNKGKEVLVLWGVHHPS    199
                     .:|||.|..|||:||:||.|||.||||:..:|:|:||||||||||:|||.
A/Victoria/48    151 AACSHAGARSFYKNLIWLVKKGKSYPKINQTYINDKGKEVLVLWGIHHPP    200

A/WSN/1933       200 SSDEQQSLYSNGNAYVSVASSNYNRRFTPEIAARPKVRDQHGRMNYYWTL    249
                     :..:|:|||.|.:|||.|.:|.|:::|.|||||||||||:.|||||||||
A/Victoria/48    201 TITDQESLYQNADAYVFVGTSRYSKKFKPEIAARPKVRDRAGRMNYYWTL    250

A/WSN/1933       250 LEPGDTIIFEATGNLIAPWYAFALSRGFESGIITSNASMHECNTKCQTPQ    299
                     :||||.|.|||||||:||.|||.:.:...||||.|:..:|:||..||||:
A/Victoria/48    251 VEPGDKITFEATGNLVAPRYAFTMEKEAGSGIIISDTPVHDCNATCQTPE    300

A/WSN/1933       300 GAINSNLPFQNIHPVTIGECPKYVRSTKLRMVTGLRNIPSIQYRGLFGAI    349
                     ||||::|||||:||:|||:|||||||||||:.|||||:||||.|||||||
A/Victoria/48    301 GAINTSLPFQNVHPITIGKCPKYVRSTKLRLATGLRNVPSIQSRGLFGAI    350

A/WSN/1933       350 AGFIEGGWTGMIDGWYGYHHQNEQGSGYAADQKSTQNAINGITNKVNSVI    399
                     |||||||||||:||||||||||:||||||||.|||||||:.|||||||||
A/Victoria/48    351 AGFIEGGWTGMVDGWYGYHHQNDQGSGYAADLKSTQNAIDKITNKVNSVI    400

A/WSN/1933       400 EKMNTQFTAVGKEFNNLEKRMENLNKKVDDGFLDIWTYNAELLVLLENER    449
                     |||||||||||||||:||||:||||||||||||||:||||||||||||||
A/Victoria/48    401 EKMNTQFTAVGKEFNHLEKRIENLNKKVDDGFLDVWTYNAELLVLLENER    450

A/WSN/1933       450 TLDFHDLNVKNLYEKVKSQLKNNAKEIGNGCFEFYHKCDNECMESVRNGT    499
                     |||:||.|||||||||:.||||||||||||||||||||||.|||||:|||
A/Victoria/48    451 TLDYHDSNVKNLYEKVRHQLKNNAKEIGNGCFEFYHKCDNTCMESVKNGT    500

A/WSN/1933       500 YDYPKYSEESKLNREKIDGVKLESMGVYQILAIYSTVASSLVLLVSLGAI    549
                     |||||||||:||||||||||||:|..:||||||||||||||||:||||||
A/Victoria/48    501 YDYPKYSEEAKLNREKIDGVKLDSTRIYQILAIYSTVASSLVLVVSLGAI    550

A/WSN/1933       550 SFWMCSNGSLQCRICI    565
                     ||||||||||||||||
A/Victoria/48    551 SFWMCSNGSLQCRICI    566
```

#### A/WSN/1933 vs A/Wisconsin/67/2022

```
A/WSN/1933          1 MKAKLLVLLYAFVATDADTICIGYHANNSTDTVDTILEKNVAVTHSVNLL     50
                      |||.|:|:||.|...:|||:|||||||||||||||:|||||.||||||||
A/Wisconsin/6       1 MKAILVVMLYTFTTANADTLCIGYHANNSTDTVDTVLEKNVTVTHSVNLL     50

A/WSN/1933         51 EDSHNGKLCKLKGIAPLQLGKCNITGWLLGNPECDSLLPARSWSYIVETP    100
                      ||.||||||||:|:|||.||:|||.||:||||||:||..||||||||||.
A/Wisconsin/6      51 EDKHNGKLCKLRGVAPLHLGQCNIAGWILGNPECESLSTARSWSYIVETS    100

A/WSN/1933        101 NSENGACYPGDLIDYEELREQLSSVSSLERFEIFPKESSWPNH-TFNGVT    149
                      ||:||.|||||.|:|||||||||||||.||||||||.|||||| :.||||
A/Wisconsin/6     101 NSDNGTCYPGDFINYEELREQLSSVSSFERFEIFPKTSSWPNHDSDNGVT    150

A/WSN/1933        150 VSCSHRGKSSFYRNLLWLTKKGDSYPKLTNSYVNNKGKEVLVLWGVHHPS    199
                      .:|||.|..|||:||:||.|||.||||:..:|:|:||||||||||:|||.
A/Wisconsin/6     151 AACSHAGARSFYKNLIWLVKKGKSYPKINQTYINDKGKEVLVLWGIHHPP    200

A/WSN/1933        200 SSDEQQSLYSNGNAYVSVASSNYNRRFTPEIAARPKVRDQHGRMNYYWTL    249
                      :..:|:|||.|.:|||.|.:|.|:::|.||||.|||||||.|||||||||
A/Wisconsin/6     201 TITDQESLYQNADAYVFVGTSRYSKKFKPEIATRPKVRDQAGRMNYYWTL    250

A/WSN/1933        250 LEPGDTIIFEATGNLIAPWYAFALSRGFESGIITSNASMHECNTKCQTPQ    299
                      :||||.|.|||||||:||.|||.:.:...||||.|:..:|:||..||||:
A/Wisconsin/6     251 VEPGDKITFEATGNLVAPRYAFTMEKEAGSGIIISDTPVHDCNATCQTPE    300

A/WSN/1933        300 GAINSNLPFQNIHPVTIGECPKYVRSTKLRMVTGLRNIPSIQYRGLFGAI    349
                      ||||::|||||:||:|||:|||||||||||:.|||||:||||.|||||||
A/Wisconsin/6     301 GAINTSLPFQNVHPITIGKCPKYVRSTKLRLATGLRNVPSIQSRGLFGAI    350

A/WSN/1933        350 AGFIEGGWTGMIDGWYGYHHQNEQGSGYAADQKSTQNAINGITNKVNSVI    399
                      |||||||||||:||||||||||:||||||||.|||||||:.|||||||||
A/Wisconsin/6     351 AGFIEGGWTGMVDGWYGYHHQNDQGSGYAADLKSTQNAIDKITNKVNSVI    400

A/WSN/1933        400 EKMNTQFTAVGKEFNNLEKRMENLNKKVDDGFLDIWTYNAELLVLLENER    449
                      |||||||||||||||:||||:|||||||||||||:|||||||||||||||
A/Wisconsin/6     401 EKMNTQFTAVGKEFNHLEKRIENLNKKVDDGFLDVWTYNAELLVLLENER    450

A/WSN/1933        450 TLDFHDLNVKNLYEKVKSQLKNNAKEIGNGCFEFYHKCDNECMESVRNGT    499
                      |||:||.|||||||||:.||||||||||||||||||||||.|||||:|||
A/Wisconsin/6     451 TLDYHDSNVKNLYEKVRHQLKNNAKEIGNGCFEFYHKCDNTCMESVKNGT    500

A/WSN/1933        500 YDYPKYSEESKLNREKIDGVKLESMGVYQILAIYSTVASSLVLLVSLGAI    549
                      |||||||||:||||||||||||:|..:||||||||||||||||:||||||
A/Wisconsin/6     501 YDYPKYSEEAKLNREKIDGVKLDSTRIYQILAIYSTVASSLVLVVSLGAI    550

A/WSN/1933        550 SFWMCSNGSLQCRICI    565
                      ||||||||||||||||
A/Wisconsin/6     551 SFWMCSNGSLQCRICI    566
```

#### A/WSN/1933 vs A/Missouri/11/2025

```
A/WSN/1933          1 MKAKLLVLLYAFVATDADTICIGYHANNSTDTVDTILEKNVAVTHSVNLL     50
                      |||.|:|:||.|...:|||:|||||||||||||||:|||||.||||||||
A/Missouri/11       1 MKAILVVMLYTFTTANADTLCIGYHANNSTDTVDTVLEKNVTVTHSVNLL     50

A/WSN/1933         51 EDSHNGKLCKLKGIAPLQLGKCNITGWLLGNPECDSLLPARSWSYIVETP    100
                      ||.||||||||:|:|||.||:|||.||:||||||:||..||||||||||.
A/Missouri/11      51 EDKHNGKLCKLRGVAPLHLGQCNIAGWILGNPECESLSTARSWSYIVETS    100

A/WSN/1933        101 NSENGACYPGDLIDYEELREQLSSVSSLERFEIFPKESSWPNH-TFNGVT    149
                      ||:||.|||||.|:|||||||||||||.||||||||.|||||| :.||||
A/Missouri/11     101 NSDNGTCYPGDFINYEELREQLSSVSSFERFEIFPKASSWPNHDSDNGVT    150

A/WSN/1933        150 VSCSHRGKSSFYRNLLWLTKKGDSYPKLTNSYVNNKGKEVLVLWGVHHPS    199
                      .:|||.|..|||:||:||.|||.||||:..:|:|:||||||||||:|||.
A/Missouri/11     151 AACSHAGARSFYKNLIWLVKKGKSYPKINQTYINDKGKEVLVLWGIHHPP    200

A/WSN/1933        200 SSDEQQSLYSNGNAYVSVASSNYNRRFTPEIAARPKVRDQHGRMNYYWTL    249
                      :..:|:|||.|.:|||.|.:|.|:::|.|||||||||||:.|||||||||
A/Missouri/11     201 TITDQESLYQNADAYVFVGTSRYSKKFKPEIAARPKVRDRAGRMNYYWTL    250
```

```
A/WSN/1933       250 LEPGDTIIFEATGNLIAPWYAFALSRGFESGIITSNASMHECNTKCQTPQ    299
                     :||||.|.|||||||:||.|||.:.:...||||.|:..:|:||..||||:
A/Missouri/11    251 VEPGDKITFEATGNLVAPRYAFTMEKEAGSGIIISDTPVHDCNATCQTPE    300

A/WSN/1933       300 GAINSNLPFQNIHPVTIGECPKYVRSTKLRMVTGLRNIPSIQYRGLFGAI    349
                     ||||::|||||:||:|||:|||||||||||:.|||||:||||.|||||||
A/Missouri/11    301 GAINTSLPFQNVHPITIGKCPKYVRSTKLRLATGLRNVPSIQSRGLFGAI    350

A/WSN/1933       350 AGFIEGGWTGMIDGWYGYHHQNEQGSGYAADQKSTQNAINGITNKVNSVI    399
                     |||||||||||:||||||||||:||||||||.||||||::.|||||||||
A/Missouri/11    351 AGFIEGGWTGMVDGWYGYHHQNDQGSGYAADLKSTQNAVDKITNKVNSVI    400

A/WSN/1933       400 EKMNTQFTAVGKEFNNLEKRMENLNKKVDDGFLDIWTYNAELLVLLENER    449
                     |||||||||||||||:||||:|||||||||||||:|||||||||||||||
A/Missouri/11    401 EKMNTQFTAVGKEFNHLEKRIENLNKKVDDGFLDVWTYNAELLVLLENER    450

A/WSN/1933       450 TLDFHDLNVKNLYEKVKSQLKNNAKEIGNGCFEFYHKCDNECMESVRNGT    499
                     |||:||.|||||||||:.||||||||.|||||||.|||||.|||||:|||
A/Missouri/11    451 TLDYHDSNVKNLYEKVRHQLKNNAKETGNGCFEFXHKCDNTCMESVKNGT    500

A/WSN/1933       500 YDYPKYSEESKLNREKIDGVKLESMGVYQILAIYSTVASSLVLLVSLGAI    549
                     |||||||||:||||||||||||:|..:|||||||||.||||||:||||||
A/Missouri/11    501 YDYPKYSEEAKLNREKIDGVKLDSTRIYQILAIYSTAASSLVLVVSLGAI    550

A/WSN/1933       550 SFWMCSNGSLQCRICI    565
                     ||||||||||||||||
A/Missouri/11    551 SFWMCSNGSLQCRICI    566
```

#### A/WSN/1933 vs A/Brisbane/59/2007

```
A/WSN/1933         1 MKAKLLVLLYAFVATDADTICIGYHANNSTDTVDTILEKNVAVTHSVNLL     50
                     ||.||||||..|.||.|||||||||||||||||||:|||||.||||||||
A/Brisbane/59      1 MKVKLLVLLCTFTATYADTICIGYHANNSTDTVDTVLEKNVTVTHSVNLL     50

A/WSN/1933        51 EDSHNGKLCKLKGIAPLQLGKCNITGWLLGNPECDSLLPARSWSYIVETP    100
                     |:|||||||.||||||||||.|::.||:||||||:.|:...|||||||.|
A/Brisbane/59     51 ENSHNGKLCLLKGIAPLQLGNCSVAGWILGNPECELLISKESWSYIVEKP    100

A/WSN/1933       101 NSENGACYPGDLIDYEELREQLSSVSSLERFEIFPKESSWPNHTFNGVTV    150
                     |.|||.||||...||||||||||||||.||||||||||||||||..||:.
A/Brisbane/59    101 NPENGTCYPGHFADYEELREQLSSVSSFERFEIFPKESSWPNHTVTGVSA    150

A/WSN/1933       151 SCSHRGKSSFYRNLLWLTKKGDSYPKLTNSYVNNKGKEVLVLWGVHHPSS    200
                     ||||.|:|||||||||||.|...||.|:.||.|||.||||||||||||.:
A/Brisbane/59    151 SCSHNGESSFYRNLLWLTGKNGLYPNLSKSYANNKEKEVLVLWGVHHPPN    200

A/WSN/1933       201 SDEQQSLYSNGNAYVSVASSNYNRRFTPEIAARPKVRDQHGRMNYYWTLL    250
                     ...|::||...||||||.||:|:|:||||||.|||||||.||:|||||||
A/Brisbane/59    201 IGIQKALYHTENAYVSVVSSHYSRKFTPEIAKRPKVRDQEGRINYYWTLL    250

A/WSN/1933       251 EPGDTIIFEATGNLIAPWYAFALSRGFESGIITSNASMHECNTKCQTPQG    300
                     ||||||||||.||||||.|||||||||.||||.|||.|.:|:.|||||||
A/Brisbane/59    251 EPGDTIIFEANGNLIAPRYAFALSRGFGSGIINSNAPMDKCDAKCQTPQG    300

A/WSN/1933       301 AINSNLPFQNIHPVTIGECPKYVRSTKLRMVTGLRNIPSIQYRGLFGAIA    350
                     ||||:|||||:||||||||||||||.|||||||||||||||.||||||||
A/Brisbane/59    301 AINSSLPFQNVHPVTIGECPKYVRSAKLRMVTGLRNIPSIQSRGLFGAIA    350

A/WSN/1933       351 GFIEGGWTGMIDGWYGYHHQNEQGSGYAADQKSTQNAINGITNKVNSVIE    400
                     ||||||||||:|||||||||||||||||||||||||||||||||||||||
A/Brisbane/59    351 GFIEGGWTGMVDGWYGYHHQNEQGSGYAADQKSTQNAINGITNKVNSVIE    400

A/WSN/1933       401 KMNTQFTAVGKEFNNLEKRMENLNKKVDDGFLDIWTYNAELLVLLENERT    450
                     ||||||||||||||.||:|||||||||||||:||||||||||||||||||
A/Brisbane/59    401 KMNTQFTAVGKEFNKLERRMENLNKKVDDGFIDIWTYNAELLVLLENERT    450

A/WSN/1933       451 LDFHDLNVKNLYEKVKSQLKNNAKEIGNGCFEFYHKCDNECMESVRNGTY    500
                     |||||.|||||||||||||||||||||||||||||||::||||||:||||
A/Brisbane/59    451 LDFHDSNVKNLYEKVKSQLKNNAKEIGNGCFEFYHKCNDECMESVKNGTY    500

A/WSN/1933       501 DYPKYSEESKLNREKIDGVKLESMGVYQILAIYSTVASSLVLLVSLGAIS    550
                     ||||||||||||||||||||||||||||||||||||||||||||||||||
A/Brisbane/59    501 DYPKYSEESKLNREKIDGVKLESMGVYQILAIYSTVASSLVLLVSLGAIS    550
```

```
A/WSN/1933       551 FWMCSNGSLQCRICI    565
                     |||||||||||||||
A/Brisbane/59    551 FWMCSNGSLQCRICI    565
```

#### A/WSN/1933 vs A/California/04/2009

```
A/WSN/1933         1 MKAKLLVLLYAFVATDADTICIGYHANNSTDTVDTILEKNVAVTHSVNLL     50
                     |||.|:||||.|...:|||:|||||||||||||||:|||||.||||||||
A/California/      1 MKAILVVLLYTFATANADTLCIGYHANNSTDTVDTVLEKNVTVTHSVNLL     50

A/WSN/1933        51 EDSHNGKLCKLKGIAPLQLGKCNITGWLLGNPECDSLLPARSWSYIVETP    100
                     ||.||||||||:|:|||.||||||.||:||||||:||..|.|||||||||
A/California/     51 EDKHNGKLCKLRGVAPLHLGKCNIAGWILGNPECESLSTASSWSYIVETP    100

A/WSN/1933       101 NSENGACYPGDLIDYEELREQLSSVSSLERFEIFPKESSWPNHTFN-GVT    149
                     :|:||.|||||.|||||||||||||||.||||||||.||||||..| |||
A/California/    101 SSDNGTCYPGDFIDYEELREQLSSVSSFERFEIFPKTSSWPNHDSNKGVT    150

A/WSN/1933       150 VSCSHRGKSSFYRNLLWLTKKGDSYPKLTNSYVNNKGKEVLVLWGVHHPS    199
                     .:|.|.|..|||:||:||.|||:|||||:.||:|:||||||||||:|||.
A/California/    151 AACPHAGAKSFYKNLIWLVKKGNSYPKLSKSYINDKGKEVLVLWGIHHPP    200

A/WSN/1933       200 SSDEQQSLYSNGNAYVSVASSNYNRRFTPEIAARPKVRDQHGRMNYYWTL    249
                     :|.:|||||.|.:.||.|.||.|:::|.||||.|||||||.|||||||||
A/California/    201 TSADQQSLYQNADTYVFVGSSRYSKKFKPEIAIRPKVRDQEGRMNYYWTL    250

A/WSN/1933       250 LEPGDTIIFEATGNLIAPWYAFALSRGFESGIITSNASMHECNTKCQTPQ    299
                     :||||.|.|||||||:.|.||||:.|...||||.|:..:|:|||.||||:
A/California/    251 VEPGDKITFEATGNLVVPRYAFAMERNAGSGIIISDTPVHDCNTTCQTPK    300

A/WSN/1933       300 GAINSNLPFQNIHPVTIGECPKYVRSTKLRMVTGLRNIPSIQYRGLFGAI    349
                     ||||::||||||||:|||:|||||:|||||:.||||||||||.|||||||
A/California/    301 GAINTSLPFQNIHPITIGKCPKYVKSTKLRLATGLRNIPSIQSRGLFGAI    350

A/WSN/1933       350 AGFIEGGWTGMIDGWYGYHHQNEQGSGYAADQKSTQNAINGITNKVNSVI    399
                     |||||||||||:|||||||||||||||||||.|||||||:.|||||||||
A/California/    351 AGFIEGGWTGMVDGWYGYHHQNEQGSGYAADLKSTQNAIDEITNKVNSVI    400

A/WSN/1933       400 EKMNTQFTAVGKEFNNLEKRMENLNKKVDDGFLDIWTYNAELLVLLENER    449
                     |||||||||||||||:||||:|||||||||||||||||||||||||||||
A/California/    401 EKMNTQFTAVGKEFNHLEKRIENLNKKVDDGFLDIWTYNAELLVLLENER    450

A/WSN/1933       450 TLDFHDLNVKNLYEKVKSQLKNNAKEIGNGCFEFYHKCDNECMESVRNGT    499
                     |||:||.|||||||||:|||||||||||||||||||||||.|||||:|||
A/California/    451 TLDYHDSNVKNLYEKVRSQLKNNAKEIGNGCFEFYHKCDNTCMESVKNGT    500

A/WSN/1933       500 YDYPKYSEESKLNREKIDGVKLESMGVYQILAIYSTVASSLVLLVSLGAI    549
                     |||||||||:|||||:||||||||..:||||||||||||||||:||||||
A/California/    501 YDYPKYSEEAKLNREEIDGVKLESTRIYQILAIYSTVASSLVLVVSLGAI    550

A/WSN/1933       550 SFWMCSNGSLQCRICI    565
                     ||||||||||||||||
A/California/    551 SFWMCSNGSLQCRICI    566
```

#### A/WSN/1933 vs A/California/07/2009

```
A/WSN/1933         1 MKAKLLVLLYAFVATDADTICIGYHANNSTDTVDTILEKNVAVTHSVNLL     50
                     |||.|:||||.|...:|||:|||||||||||||||:|||||.||||||||
A/California/      1 MKAILVVLLYTFATANADTLCIGYHANNSTDTVDTVLEKNVTVTHSVNLL     50

A/WSN/1933        51 EDSHNGKLCKLKGIAPLQLGKCNITGWLLGNPECDSLLPARSWSYIVETP    100
                     ||.||||||||:|:|||.||||||.||:||||||:||..|.|||||||||
A/California/     51 EDKHNGKLCKLRGVAPLHLGKCNIAGWILGNPECESLSTASSWSYIVETP    100

A/WSN/1933       101 NSENGACYPGDLIDYEELREQLSSVSSLERFEIFPKESSWPNHTFN-GVT    149
                     :|:||.|||||.|||||||||||||||.||||||||.||||||..| |||
A/California/    101 SSDNGTCYPGDFIDYEELREQLSSVSSFERFEIFPKTSSWPNHDSNKGVT    150

A/WSN/1933       150 VSCSHRGKSSFYRNLLWLTKKGDSYPKLTNSYVNNKGKEVLVLWGVHHPS    199
                     .:|.|.|..|||:||:||.|||:|||||:.||:|:||||||||||:||||
A/California/    151 AACPHAGAKSFYKNLIWLVKKGNSYPKLSKSYINDKGKEVLVLWGIHHPS    200
```

```
A/WSN/1933       200 SSDEQQSLYSNGNAYVSVASSNYNRRFTPEIAARPKVRDQHGRMNYYWTL    249
                     :|.:|||||.|.:|||.|.||.|:::|.||||.|||||...|||||||||
A/California/    201 TSADQQSLYQNADAYVFVGSSRYSKKFKPEIAIRPKVRXXEGRMNYYWTL    250

A/WSN/1933       250 LEPGDTIIFEATGNLIAPWYAFALSRGFESGIITSNASMHECNTKCQTPQ    299
                     :||||.|.|||||||:.|.||||:.|...||||.|:..:|:|||.||||:
A/California/    251 VEPGDKITFEATGNLVVPRYAFAMERNAGSGIIISDTPVHDCNTTCQTPK    300

A/WSN/1933       300 GAINSNLPFQNIHPVTIGECPKYVRSTKLRMVTGLRNIPSIQYRGLFGAI    349
                     ||||::||||||||:|||:|||||:|||||:.||||||||||.|||||||
A/California/    301 GAINTSLPFQNIHPITIGKCPKYVKSTKLRLATGLRNIPSIQSRGLFGAI    350

A/WSN/1933       350 AGFIEGGWTGMIDGWYGYHHQNEQGSGYAADQKSTQNAINGITNKVNSVI    399
                     |||||||||||:|||||||||||||||||||.|||||||:.|||||||||
A/California/    351 AGFIEGGWTGMVDGWYGYHHQNEQGSGYAADLKSTQNAIDEITNKVNSVI    400

A/WSN/1933       400 EKMNTQFTAVGKEFNNLEKRMENLNKKVDDGFLDIWTYNAELLVLLENER    449
                     |||||||||||||||:||||:|||||||||||||||||||||||||||||
A/California/    401 EKMNTQFTAVGKEFNHLEKRIENLNKKVDDGFLDIWTYNAELLVLLENER    450

A/WSN/1933       450 TLDFHDLNVKNLYEKVKSQLKNNAKEIGNGCFEFYHKCDNECMESVRNGT    499
                     |||:||.|||||||||:|||||||||||||||||||||||.|||||:|||
A/California/    451 TLDYHDSNVKNLYEKVRSQLKNNAKEIGNGCFEFYHKCDNTCMESVKNGT    500

A/WSN/1933       500 YDYPKYSEESKLNREKIDGVKLESMGVYQILAIYSTVASSLVLLVSLGAI    549
                     |||||||||:|||||:||||||||..:||||||||||||||||:||||||
A/California/    501 YDYPKYSEEAKLNREEIDGVKLESTRIYQILAIYSTVASSLVLVVSLGAI    550

A/WSN/1933       550 SFWMCSNGSLQCRICI    565
                     ||||||||||||||||
A/California/    551 SFWMCSNGSLQCRICI    566
```

#### A/WSN/1933 vs A/Moscow/IIV01/2009

```
A/WSN/1933         1 MKAKLLVLLYAFVATDADTICIGYHANNSTDTVDTILEKNVAVTHSVNLL     50
                     |||.|:||||.|...:|||:|||||||||||||||:|||||.||||||||
A/Moscow/IIV0      1 MKAILVVLLYTFATANADTLCIGYHANNSTDTVDTVLEKNVTVTHSVNLL     50

A/WSN/1933        51 EDSHNGKLCKLKGIAPLQLGKCNITGWLLGNPECDSLLPARSWSYIVETP    100
                     ||.||||||||:|:|||.||||||.||:||||||:||..|.||||||||.
A/Moscow/IIV0     51 EDKHNGKLCKLRGVAPLHLGKCNIAGWILGNPECESLSTASSWSYIVETS    100

A/WSN/1933       101 NSENGACYPGDLIDYEELREQLSSVSSLERFEIFPKESSWPNHTFN-GVT    149
                     :|:||.|||||.|||||||||||||||.||||||||.||||||..| |||
A/Moscow/IIV0    101 SSDNGTCYPGDFIDYEELREQLSSVSSFERFEIFPKTSSWPNHDSNKGVT    150

A/WSN/1933       150 VSCSHRGKSSFYRNLLWLTKKGDSYPKLTNSYVNNKGKEVLVLWGVHHPS    199
                     .:|.|.|..|||:||:||.|||:|||||:.||:|:||||||||||:||||
A/Moscow/IIV0    151 AACPHAGAKSFYKNLIWLVKKGNSYPKLSKSYINDKGKEVLVLWGIHHPS    200

A/WSN/1933       200 SSDEQQSLYSNGNAYVSVASSNYNRRFTPEIAARPKVRDQHGRMNYYWTL    249
                     :|.:|||||.|.:|||.|.:|.|:::|.||||.|||||||.|||||||||
A/Moscow/IIV0    201 TSADQQSLYQNADAYVFVGTSRYSKKFKPEIAIRPKVRDQEGRMNYYWTL    250

A/WSN/1933       250 LEPGDTIIFEATGNLIAPWYAFALSRGFESGIITSNASMHECNTKCQTPQ    299
                     :||||.|.|||||||:.|.||||:.|...||||.|:..:|:|||.||||:
A/Moscow/IIV0    251 VEPGDKITFEATGNLVVPRYAFAMERNAGSGIIISDTPVHDCNTTCQTPK    300

A/WSN/1933       300 GAINSNLPFQNIHPVTIGECPKYVRSTKLRMVTGLRNIPSIQYRGLFGAI    349
                     ||||::||||||||:|||:|||||:|||||:.|||||:||||.|||||||
A/Moscow/IIV0    301 GAINTSLPFQNIHPITIGKCPKYVKSTKLRLATGLRNVPSIQSRGLFGAI    350

A/WSN/1933       350 AGFIEGGWTGMIDGWYGYHHQNEQGSGYAADQKSTQNAINGITNKVNSVI    399
                     |||||||||||:|||||||||||||||||||.|||||||:.|||||||||
A/Moscow/IIV0    351 AGFIEGGWTGMVDGWYGYHHQNEQGSGYAADLKSTQNAIDEITNKVNSVI    400

A/WSN/1933       400 EKMNTQFTAVGKEFNNLEKRMENLNKKVDDGFLDIWTYNAELLVLLENER    449
                     |||||||||||||||:||||:|||||||||||||||||||||||||||||
A/Moscow/IIV0    401 EKMNTQFTAVGKEFNHLEKRIENLNKKVDDGFLDIWTYNAELLVLLENER    450

A/WSN/1933       450 TLDFHDLNVKNLYEKVKSQLKNNAKEIGNGCFEFYHKCDNECMESVRNGT    499
                     |||:||.|||||||||:|||||||||||||||||||||||.|||||:|||
A/Moscow/IIV0    451 TLDYHDSNVKNLYEKVRSQLKNNAKEIGNGCFEFYHKCDNTCMESVKNGT    500
```

```
A/WSN/1933       500 YDYPKYSEESKLNREKIDGVKLESMGVYQILAIYSTVASSLVLLVSLGAI    549
                     |||||||||:|||||:||||||||..:||||||||||||||||:||||||
A/Moscow/IIV0    501 YDYPKYSEEAKLNREEIDGVKLESTRIYQILAIYSTVASSLVLVVSLGAI    550

A/WSN/1933       550 SFWMCSNGSLQCRICI    565
                     ||||||||||||||||
A/Moscow/IIV0    551 SFWMCSNGSLQCRICI    566
```

#### A/WSN/1933 vs A/Narita/1/2009

```
A/WSN/1933         1 MKAKLLVLLYAFVATDADTICIGYHANNSTDTVDTILEKNVAVTHSVNLL     50
                     |||.|:||||.|...:|||:|||||||||||||||:|||||.||||||||
A/Narita/1/20      1 MKAILVVLLYTFATANADTLCIGYHANNSTDTVDTVLEKNVTVTHSVNLL     50

A/WSN/1933        51 EDSHNGKLCKLKGIAPLQLGKCNITGWLLGNPECDSLLPARSWSYIVETP    100
                     ||.||||||||:|:|||.||||||.||:||||||:||..|.||||||||.
A/Narita/1/20     51 EDKHNGKLCKLRGVAPLHLGKCNIAGWILGNPECESLSTASSWSYIVETS    100

A/WSN/1933       101 NSENGACYPGDLIDYEELREQLSSVSSLERFEIFPKESSWPNHTFN-GVT    149
                     :|:||.|||||.|||||||||||||||.||||||||.||||||..| |||
A/Narita/1/20    101 SSDNGTCYPGDFIDYEELREQLSSVSSFERFEIFPKTSSWPNHDSNKGVT    150

A/WSN/1933       150 VSCSHRGKSSFYRNLLWLTKKGDSYPKLTNSYVNNKGKEVLVLWGVHHPS    199
                     .:|.|.|..|||:||:||.|||:|||||:.||:|:||||||||||:||||
A/Narita/1/20    151 AACPHAGAKSFYKNLIWLVKKGNSYPKLSKSYINDKGKEVLVLWGIHHPS    200

A/WSN/1933       200 SSDEQQSLYSNGNAYVSVASSNYNRRFTPEIAARPKVRDQHGRMNYYWTL    249
                     :|.:|||||.|.:|||.|.||.|:::|.||||.|||||||.|||||||||
A/Narita/1/20    201 TSADQQSLYQNADAYVFVGSSRYSKKFKPEIAIRPKVRDQEGRMNYYWTL    250

A/WSN/1933       250 LEPGDTIIFEATGNLIAPWYAFALSRGFESGIITSNASMHECNTKCQTPQ    299
                     :||||.|.|||||||:.|.||||:.|...||||.|:..:|:|||.||||:
A/Narita/1/20    251 VEPGDKITFEATGNLVVPRYAFAMERNAGSGIIISDTPVHDCNTTCQTPK    300

A/WSN/1933       300 GAINSNLPFQNIHPVTIGECPKYVRSTKLRMVTGLRNIPSIQYRGLFGAI    349
                     ||||::||||||||:|||:|||||:|||||:.|||||:||||.|||||||
A/Narita/1/20    301 GAINTSLPFQNIHPITIGKCPKYVKSTKLRLATGLRNVPSIQSRGLFGAI    350

A/WSN/1933       350 AGFIEGGWTGMIDGWYGYHHQNEQGSGYAADQKSTQNAINGITNKVNSVI    399
                     |||||||||||:|||||||||||||||||||.|||||||:.|||||||||
A/Narita/1/20    351 AGFIEGGWTGMVDGWYGYHHQNEQGSGYAADLKSTQNAIDEITNKVNSVI    400

A/WSN/1933       400 EKMNTQFTAVGKEFNNLEKRMENLNKKVDDGFLDIWTYNAELLVLLENER    449
                     |||||||||||||||:||||:|||||||||||||||||||||||||||||
A/Narita/1/20    401 EKMNTQFTAVGKEFNHLEKRIENLNKKVDDGFLDIWTYNAELLVLLENER    450

A/WSN/1933       450 TLDFHDLNVKNLYEKVKSQLKNNAKEIGNGCFEFYHKCDNECMESVRNGT    499
                     |||:||.|||||||||:|||||||||||||||||||||||.|||||:|||
A/Narita/1/20    451 TLDYHDSNVKNLYEKVRSQLKNNAKEIGNGCFEFYHKCDNTCMESVKNGT    500

A/WSN/1933       500 YDYPKYSEESKLNREKIDGVKLESMGVYQILAIYSTVASSLVLLVSLGAI    549
                     |||||||||:|||||:||||||||..:||||||||||||||||:||||||
A/Narita/1/20    501 YDYPKYSEEAKLNREEIDGVKLESTRIYQILAIYSTVASSLVLVVSLGAI    550

A/WSN/1933       550 SFWMCSNGSLQCRICI    565
                     ||||||||||||||||
A/Narita/1/20    551 SFWMCSNGSLQCRICI    566
```

#### A/WSN/1933 vs A/Netherlands/602/2009

```
A/WSN/1933         1 MKAKLLVLLYAFVATDADTICIGYHANNSTDTVDTILEKNVAVTHSVNLL     50
                     |||.|:||||.|...:|||:|||||||||||||||:|||||.||||||||
A/Netherlands      1 MKAILVVLLYTFATANADTLCIGYHANNSTDTVDTVLEKNVTVTHSVNLL     50

A/WSN/1933        51 EDSHNGKLCKLKGIAPLQLGKCNITGWLLGNPECDSLLPARSWSYIVETP    100
                     ||.||||||||:|:|||.||||||.||:||||||:||..|.||||||||.
A/Netherlands     51 EDKHNGKLCKLRGVAPLHLGKCNIAGWILGNPECESLSTASSWSYIVETS    100
```

```
A/WSN/1933        101 NSENGACYPGDLIDYEELREQLSSVSSLERFEIFPKESSWPNHTFN-GVT    149
                      :|:||.|||||.|||||||||||||||.||||||||.||||||..| |||
A/Netherlands     101 SSDNGTCYPGDFIDYEELREQLSSVSSFERFEIFPKTSSWPNHDSNKGVT    150

A/WSN/1933        150 VSCSHRGKSSFYRNLLWLTKKGDSYPKLTNSYVNNKGKEVLVLWGVHHPS    199
                      .:|.|.|..|||:||:||.|||||||||:.||:|:||||||||||:||||
A/Netherlands     151 AACPHAGAKSFYKNLIWLVKKGDSYPKLSKSYINDKGKEVLVLWGIHHPS    200

A/WSN/1933        200 SSDEQQSLYSNGNAYVSVASSNYNRRFTPEIAARPKVRDQHGRMNYYWTL    249
                      :|.:|||||.|.:|||.|.||.|:::|.||||.|||||||.|||||||||
A/Netherlands     201 TSADQQSLYQNADAYVFVGSSRYSKKFKPEIAIRPKVRDQEGRMNYYWTL    250

A/WSN/1933        250 LEPGDTIIFEATGNLIAPWYAFALSRGFESGIITSNASMHECNTKCQTPQ    299
                      :||||.|.|||||||:.|.||||:.|...||||.|:..:|:|||.||||:
A/Netherlands     251 VEPGDKITFEATGNLVVPRYAFAMERNAGSGIIISDTPVHDCNTTCQTPK    300

A/WSN/1933        300 GAINSNLPFQNIHPVTIGECPKYVRSTKLRMVTGLRNIPSIQYRGLFGAI    349
                      ||||::||||||||:|||:|||||:|||||:.|||||:||||.|||||||
A/Netherlands     301 GAINTSLPFQNIHPITIGKCPKYVKSTKLRLATGLRNVPSIQSRGLFGAI    350

A/WSN/1933        350 AGFIEGGWTGMIDGWYGYHHQNEQGSGYAADQKSTQNAINGITNKVNSVI    399
                      |||||||||||:|||||||||||||||||||.|||||||:.|||||||||
A/Netherlands     351 AGFIEGGWTGMVDGWYGYHHQNEQGSGYAADLKSTQNAIDEITNKVNSVI    400

A/WSN/1933        400 EKMNTQFTAVGKEFNNLEKRMENLNKKVDDGFLDIWTYNAELLVLLENER    449
                      |||||||||||||||:||||:|||||||||||||||||||||||||||||
A/Netherlands     401 EKMNTQFTAVGKEFNHLEKRIENLNKKVDDGFLDIWTYNAELLVLLENER    450

A/WSN/1933        450 TLDFHDLNVKNLYEKVKSQLKNNAKEIGNGCFEFYHKCDNECMESVRNGT    499
                      |||:||.|||||||||:|||||||||||||||||||||||.|||||:|||
A/Netherlands     451 TLDYHDSNVKNLYEKVRSQLKNNAKEIGNGCFEFYHKCDNTCMESVKNGT    500

A/WSN/1933        500 YDYPKYSEESKLNREKIDGVKLESMGVYQILAIYSTVASSLVLLVSLGAI    549
                      |||||||||:|||||:||||||||..:||||||||||||||||:||||||
A/Netherlands     501 YDYPKYSEEAKLNREEIDGVKLESTRIYQILAIYSTVASSLVLVVSLGAI    550

A/WSN/1933        550 SFWMCSNGSLQCRICI    565
                      ||||||||||||||||
A/Netherlands     551 SFWMCSNGSLQCRICI    566
```

**Figure S6. Amino acid sequence alignments of hemagglutinin from several natural strains with hemagglutinin from the A/WSN/1933 strain.** Alignments were performed using EMBOSS Needle from the EMBL-EBI with the Pairwise Sequence Alignment (PSA) option. Default parameters were used, including the pairs output format and the BLOSUM62 matrix.

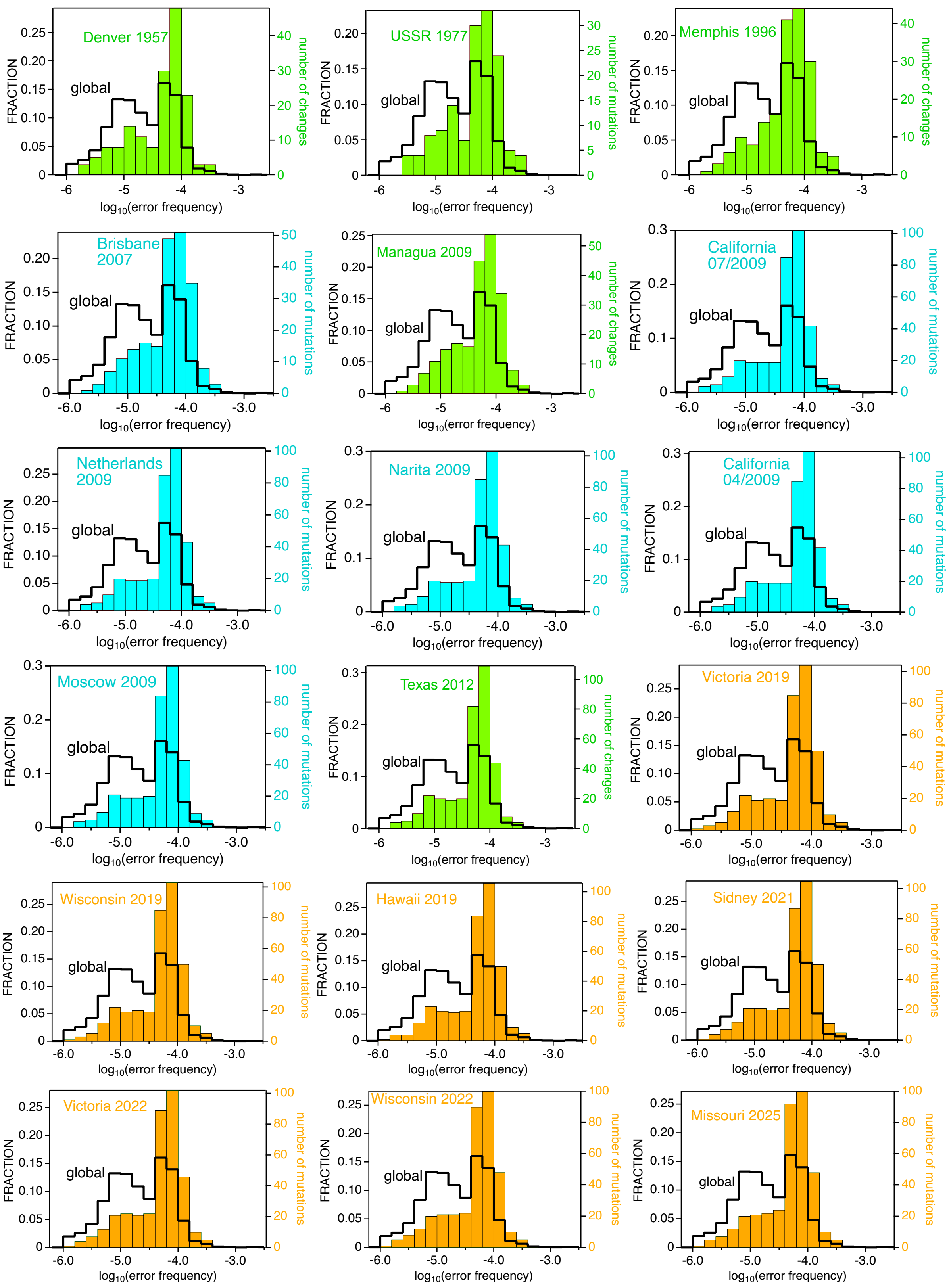


**Figure S7. Error frequency distributions for the amino acid sequence differences between hemagglutinin from natural H1 influenza strain and hemagglutinin from A/WSN/1933.** For comparison, the global distribution of Figure 2B is shown here in a cityscape representation. The color code refers to the three strain sets analyzed (see main text for details): green (set-1), orange (set-2) and blue (set-3).

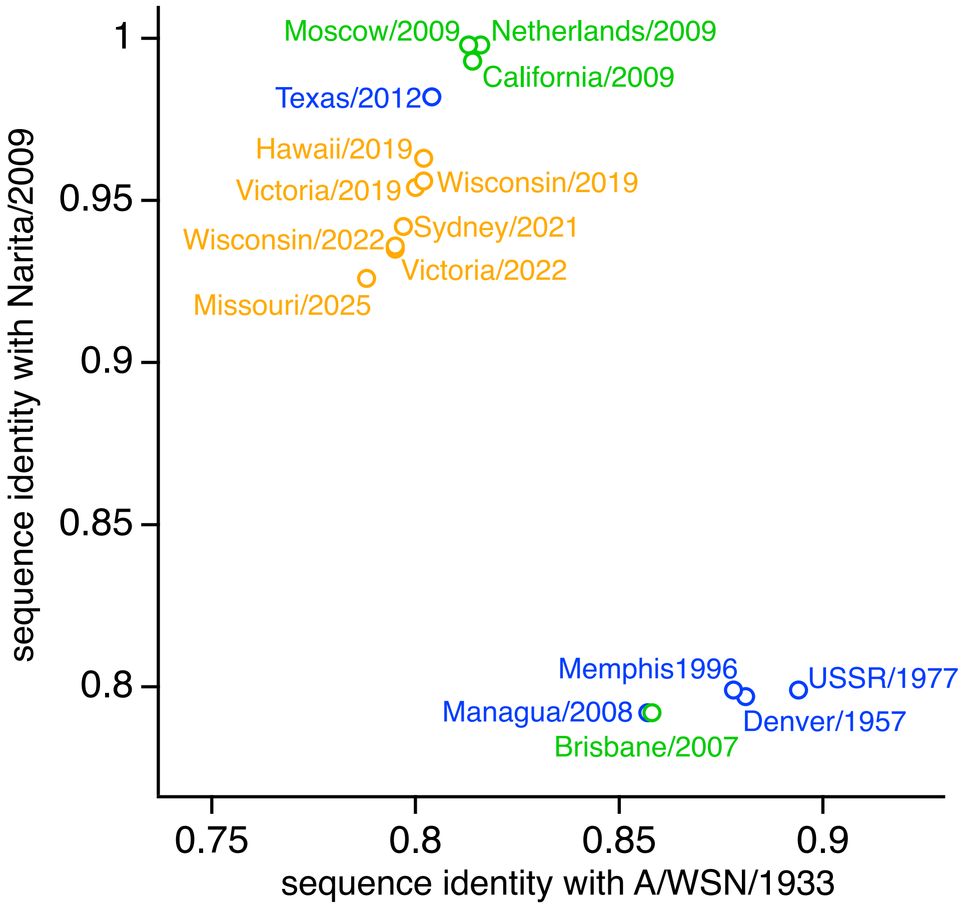


**Figure S8. Identity of hemagglutinin sequences from several natural strains with the hemagglutinins from Narita/2009 and A/WSN/1933.** Identity values are calculated from pairwise alignments of the amino acid sequences. The color code refers to the three strain sets analyzed (see main text for details): green (set-1), orange (set-2) and blue (set-3). Narita/2009 has been selected for this calculation as a representative of the viruses related to the pandemic A/H1N1pdm09 virus that emerged in 2009 in the United States and Mexico (Smith et al., 2009). Actually, Narita/2009 has been reported to be the first isolate of the pdm09 virus in Japan (Matzusaki et al., 2014). The plot of hemagglutinin identity with Narita/2009 vs. identity with A/WSN/1933 reveals that, except for Brisbane/2007, all the hemagglutinins from strains in set-3 (green) are closely related to the hemagglutinin of the pandemic A/H1N1pdm09. Furthermore, the hemagglutinins for set-2 strains (orange: strains recommended by the World Health Organization for inclusion in vaccines for the influenza seasons spanning the period 2019-2025) appear clearly to have evolved from hemagglutinin from pdm09 viruses.

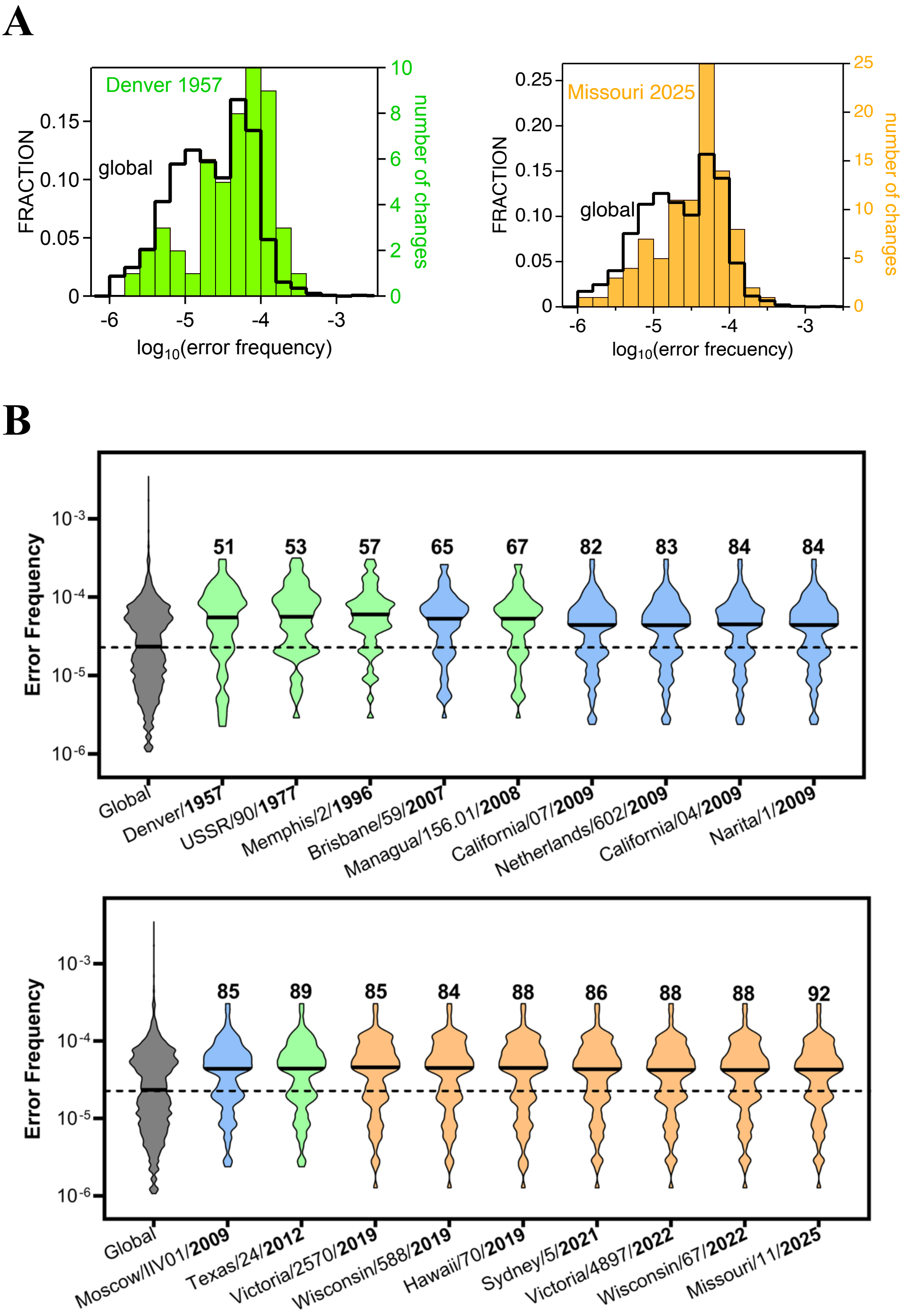


**Figure S9. Sampling of the hemagglutinin error landscape during natural influenza evolution at the amino acid sequence level.** (A) Error frequency distributions for the amino acid sequence differences between hemagglutinin from natural H1 influenza strain and hemagglutinin from A/WSN/1933. Decimal logarithms of the frequencies are binned, and bin size is 0.2. global distribution of Figure 2B is shown here in a cityscape representation. Only two illustrative examples are shown here, but all the calculated distributions are provided in panel B as violin plots. (B) Error frequency distributions for the amino acid sequence differences between hemagglutinin from natural H1 influenza strain and hemagglutinin from A/WSN/1933 shown as violin plots. The number of amino acid differences used for each strain is shown as a number above the violin. The color code refers to the three strain sets analyzed: green (set-1), orange (set-2) and blue (set-3). The sets are described in the main text, but we note here that set-2 (orange) includes strains recommended by the World Health Organization for inclusion in vaccines for the influenza seasons spanning the period 2019-2025. For comparison the violin plot for the global distribution of Figure 2B is shown in grey. In all cases, the thick horizontal segment represents the median of the distribution. To facilitate global comparison, the median segment for the global (grey) violin has been extended as a dashed line.

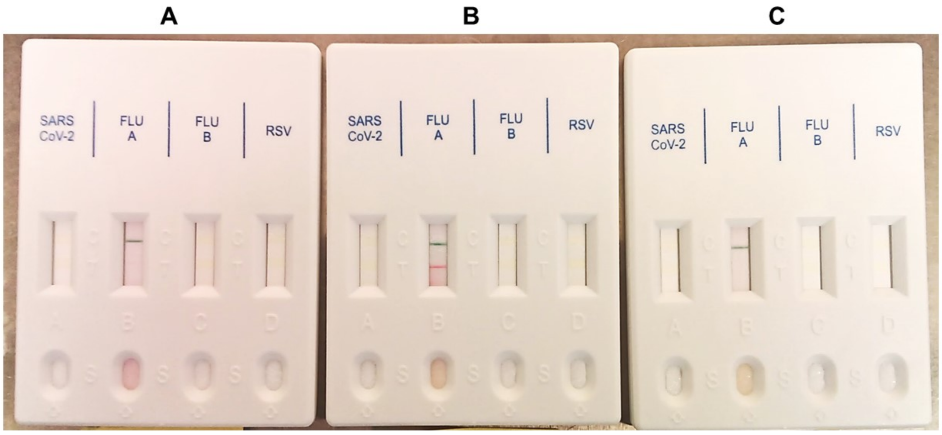


**Figure S10. Antigen-tests of influenza A from infected or mock-infected cells supernatants.** (A) 6 hours post-infection (hpi) infected cells supernatant. Negative test because of the lack of new virions in the supernatant in the first hours of infection. (B) 72 hpi infected cells supernatant. Positive test because of the presence of new virions after several days of infection (end point of the experiment). (C) 72 hpi mock-infected cells supernatant. Negative test as control of no-infection.

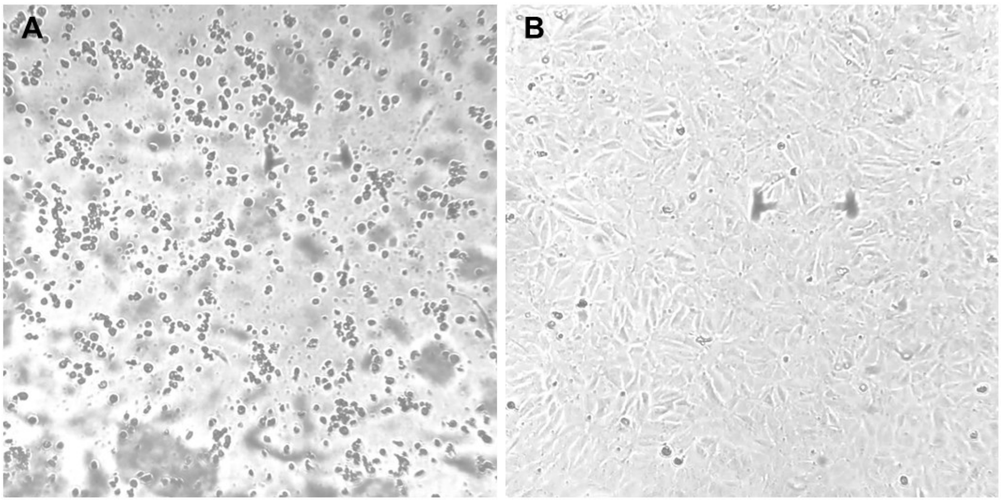


**Figure S11. Optical microscope images (400x) of infected (A) and mock-infected (B) MDCK cells, after 72 hours post-infection (hpi).** In panel A, dead adherent-cells floating in the supernatant. In panel B, alive adherent-cells remain adhered in the surface of the culture flask. Images modified to increase contrast and improve viewing.

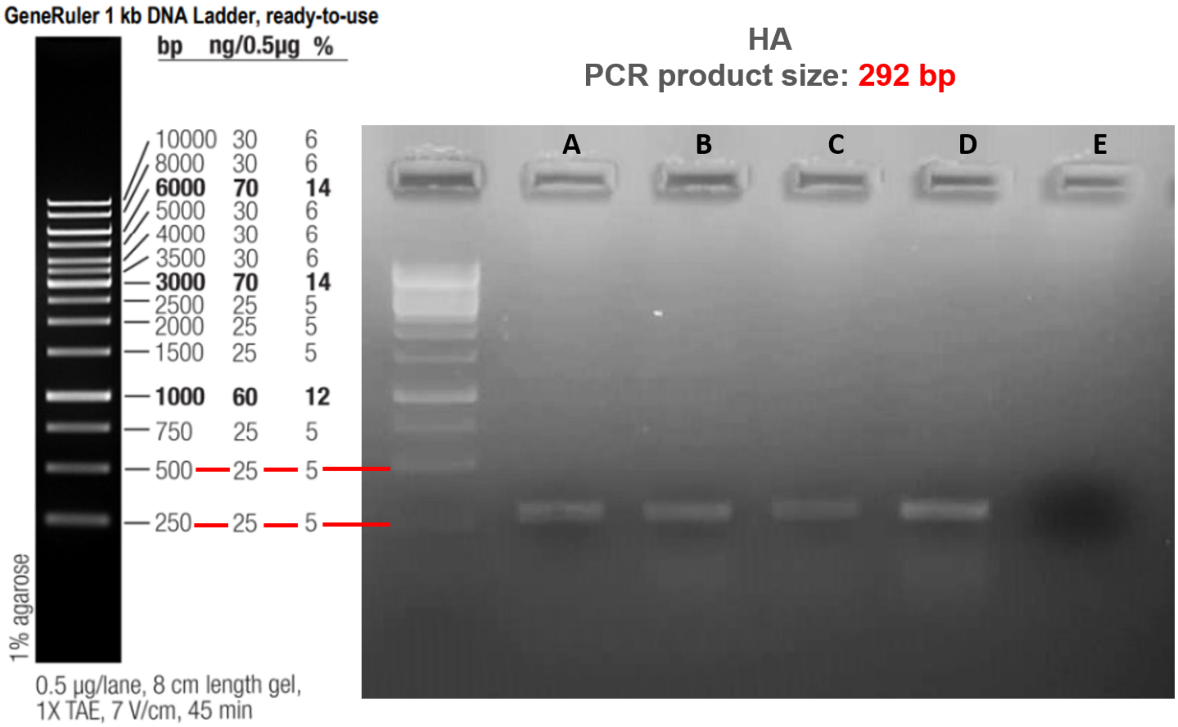


**Figure S12. Agarose gel Eelectrophoresis (2%) showing amplification of a 292 bp size PCR product of influenza A virus hemagglutinin gene.** PCR carried out on different samples: A) RNA extracted from virions (vRNA) obtained 72 hours post-infection (supernatant isolation); B), C), D) intracellular RNA from infected MDCK cells, in which hemagglutinin-encoding mRNA is expected, at 2, 4 and 6 hours post-infection, respectively; E) PCR negative control.
